# RWRtoolkit: multi-omic network analysis using random walks on multiplex networks in any species

**DOI:** 10.1101/2024.07.17.603975

**Authors:** David Kainer, Matthew Lane, Kyle A. Sullivan, J. Izaak Miller, Mikaela Cashman, Mallory Morgan, Ashley Cliff, Jonathon Romero, Angelica Walker, D. Dakota Blair, Hari Chhetri, Yongqin Wang, Mirko Pavicic, Anna Furches, Jaclyn Noshay, Meghan Drake, Natalie Landry, AJ Ireland, Ali Missaoui, Yun Kang, John Sedbrook, Paramvir Dehal, Shane Canon, Daniel Jacobson

## Abstract

Leveraging the use of multiplex multi-omic networks, key insights into genetic and epigenetic mechanisms supporting biofuel production have been uncovered. Here, we introduce RWRtoolkit, a multiplex generation, exploration, and statistical package built for R and command line users. RWRtoolkit enables the efficient exploration of large and highly complex biological networks generated from custom experimental data and/or from publicly available datasets, and is species agnostic. A range of functions can be used to find topological distances between biological entities, determine relationships within sets of interest, search for topological context around sets of interest, and statistically evaluate the strength of relationships within and between sets. The command-line interface is designed for parallelisation on high performance cluster systems, which enables high throughput analysis such as permutation testing. Several tools in the package have also been made available for use in reproducible workflows via the KBase web application.

## 1. Background

Biological studies are increasingly pursuing and obtaining data on larger scales and at multiple levels in the molecular hierarchy of the study system. One approach for dealing with the multiplicity of data in modern biology is to represent the relationships in the data as a network [1]. Each entity in a dataset (e.g., each gene) becomes a node, and an edge between two nodes represents a relationship that has been measured or predicted between those nodes (e.g., their co-expression in a population, sharing of common protein domains, similarity of methylation state, etc.). Once in network form, a great variety of network analysis methods become available [2,3]. Genes that are strongly connected to each other are topologically more likely to be functionally relevant to each other than more distal or loosely connected genes in the network. Machine learning algorithms can be used to efficiently explore entire networks and find such relationships with respect to a set of starting genes, often called seeds or anchors. This approach is particularly useful for exploring the functional context around sets of genes, such as those produced from GWAS, QTL mapping, differential expression analysis or case/control proteomics.

To analyze a set of genes in a network context, one requires the underlying network itself, and an algorithm that traverses that network. The underlying network may be as simple as a single-layer of nodes and edges generated from one experimental dataset that predicts relationships between genes, such as co/predictive expression relationship determined from RNA-seq results [4,5]. If further data types are available, (e.g., Protein-Protein Interactions [PPI]) a multi-layer network can be generated where edges in each layer represent different types of relationships between genes. A multi-layer network potentially fills in relationship gaps that exist in any given single layer, and enables simultaneous exploration of multi-omic data since one can traverse the network from gene to gene using edges from all layers. Edge weights, if they exist, can be used to influence the traversal. When relationships between specific genes are present in multiple layers, a multi-layer network presents multiple lines-of-evidence (LOE) that those genes are functionally related [6].

Commonly the multiple layers of input networks are aggregated into a single layer by summing or averaging multiple edges between the same pair of nodes into one composite edge. The result is a single adjacency matrix representation of the data, also known as a monoplex network. The aggregated network is a summarization of the input layers, and as such has lost the unique topological information carried by each layer. A more recent innovation, known as Multiplex networks, maintains the topological separation of each input layer through the use of a supra-adjacency matrix (**Fig. 1**), while still allowing for simultaneous exploration of all layers. The predictive ability of a multiplex network is often greater than the equivalent aggregated monoplex network [7,8], but the supra-adjacency matrix is more difficult to build and adds an extra level of complexity to network exploration algorithms.

**Figure 1.**
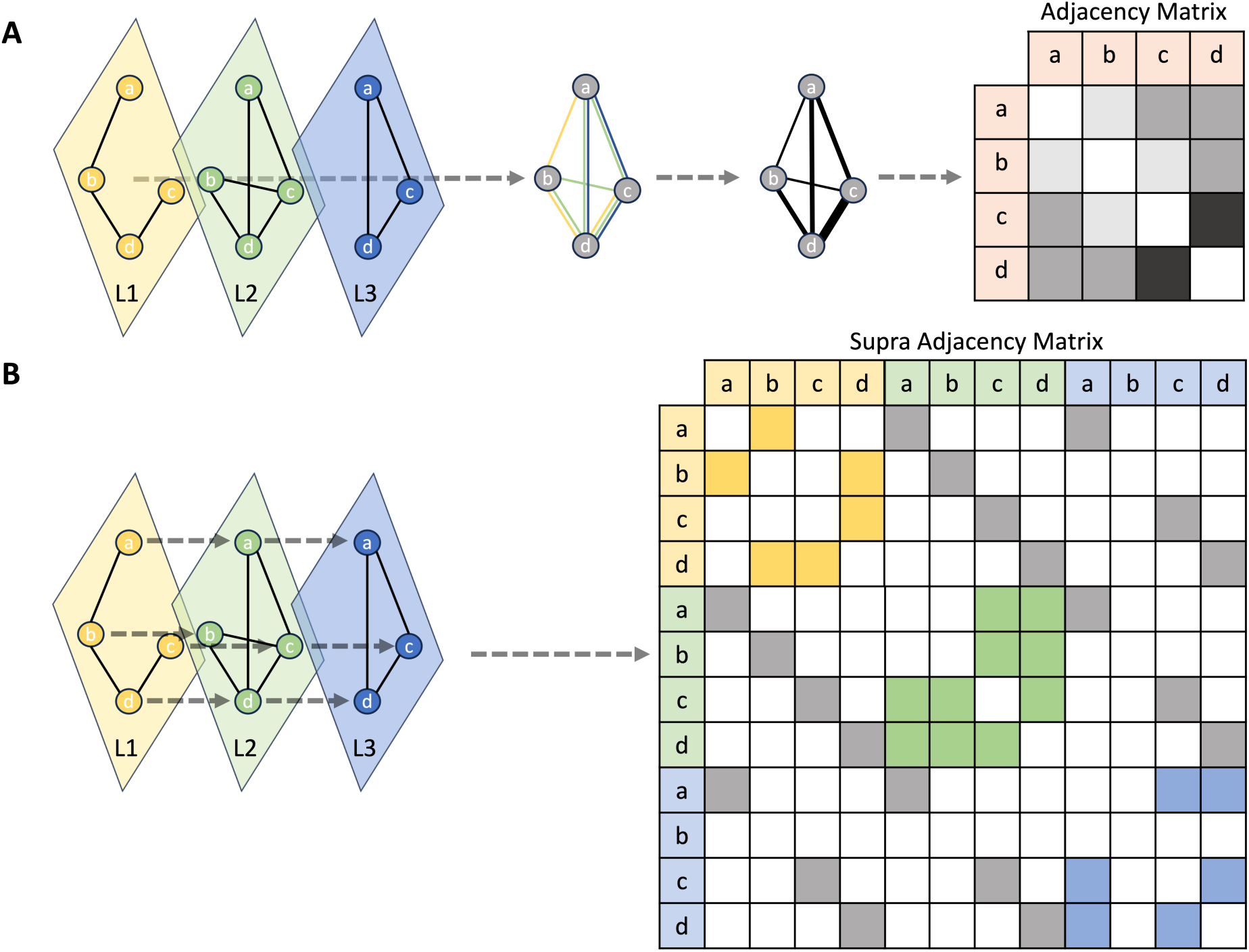
Aggregated monoplex network vs Multiplex network. In this example there are three small input network layers (L1, L2, L3), with a union set of nodes of size n=4, with which to generate a multi-layer network. **A.** In the aggregated approach the layers are merged into one. If multiple edges occur between any pair of nodes, their weights are aggregated to produce the final adjacency matrix of size n x n. **B.** In the multiplex approach each layer is kept separate via a supra-adjacency matrix of size (n x L) X (n x L) where L is the number of layers. Nodes that are common across layers are connected by virtual edges (red arrows). The diagonal blocks of the supra-adjacency matrix represent the standard adjacency matrices within each individual layer. Connectivity between layers is represented in the off-diagonal blocks, with virtual edges coloured in gray. Note that layer L3 (blue) does not contain node ‘b’, so there are no inter-layer virtual edges from L1-L3 or L2-L3 for node ‘b’.

A wide variety of algorithms exist for ranking genes according to their topological connectivity to the seed (candidate) genes in the underlying network. One simple approach, known as neighbor voting[9], scores each gene by counting their outgoing edges that directly connect to the seeds. However, by only looking at immediate connections to the seeds, the influence and importance of genes farther away is ignored. More advanced propagation approaches, such as diffusion and random walk with restart (RWR) use the entire network topology to score and rank every node, and have been shown to be generally superior in their ability to find true positive relationships [10–12]. A Random Walk can be described conceptually as a “walker” which proceeds to wander outwards from a starting seed gene, choosing which edge to take with a probability equal to 1/*d* where *d* is the degree of the current gene (*d* is the out-degree for directed networks). Over multiple iterations, the walker explores the network in this manner so the proportion of time spent at each gene forms a probability distribution that represents how accessible every gene in the network is when starting from the seed gene(s). With RWR, at each iteration the walker teleports back to the starting point with restart probability *r* to prevent the walker from wandering too far in the global topology, or getting stuck in various topological structures.

Here we introduce RWRtoolkit, an R package with a corresponding set of command line tools that enables the easy construction of multiplex networks from any set of data layers, followed by analysis of candidate gene sets within the networks using the Random Walk with Restart (RWR) algorithm. RWRtoolkit wraps the RandomWalkRestartMH R package [8], which provides the core functionality to generate multiplex networks from a set of input network layers, and implements the Random Walk Restart algorithm on a supra-adjacency matrix. Once a multiplex network has been generated, the RWRtoolkit provides commands to rank all genes in the overall network according to their connectivity to a set of seed genes, use cross-validation to assess the network’s predictive ability or determine the functional similarity of a set of genes, and find shortest paths between sets of seed genes. RWRtoolkit provides as output detailed tables of ranked genes as well as statistics of predictive accuracy (AUROC, AUPRC, etc.), plots, and network visualizations of the multi-omic neighborhood around the seed genes. Furthermore, RWRtoolkit commands can be run from the command-line interface, which enables high throughput parallelised analysis (such as permutation testing) on compute clusters.

To date, multi-omic networks have been made publicly available in a range of model species: AraNet, PopGenie, StringDB, YeastNet. However these networks are often aggregated into a single layer rather than multiplexed, and it is difficult or impossible for the user to customize their choice of input layers or include custom layers generated from their own experimental results or algorithms. RWRtoolkit enables robust network analysis of multi-omic data for any species, allowing researchers to use their own datasets and networks and/or pre-existing networks. We demonstrate this by generating a custom *Arabidopsis thaliana* multiplex network and using it to analyze gene sets from a novel GWAS study and a published gene knockout study [13]. RWRtoolkit is available with installation instructions, user guide and sample data at http://github.com/dkainer/RWRtoolkit [14]. Finally, RWRtoolkit, together with a comprehensive Arabidopsis multiplex network, has been integrated into the US Department of Energy’s KBase platform that facilitates reproducible workflows for biological analysis [15].

## 2. Data Description

We created the RWRtoolkit R package as a collection of functions for easy network-based functional analysis using random walks on multiplex networks (**Fig. 2**). The package provides functions for multiplex network generation, running random walks starting from given seed sets, validation functions for seed sets as well as network layers, and general multiplex network statistics. These functions are made available via a cross-platform R package and command-line interface commands. We have provided multiple tutorials in the R vignette format for users to explore at their own leisure. Our methods were generated with gene relationships as a focus, but these random walk methods can be applicable to any data type in network format. Example methods can be seen in Table 1 and example input and output can be seen in supplementary tables 1-18.

**Figure 2.**
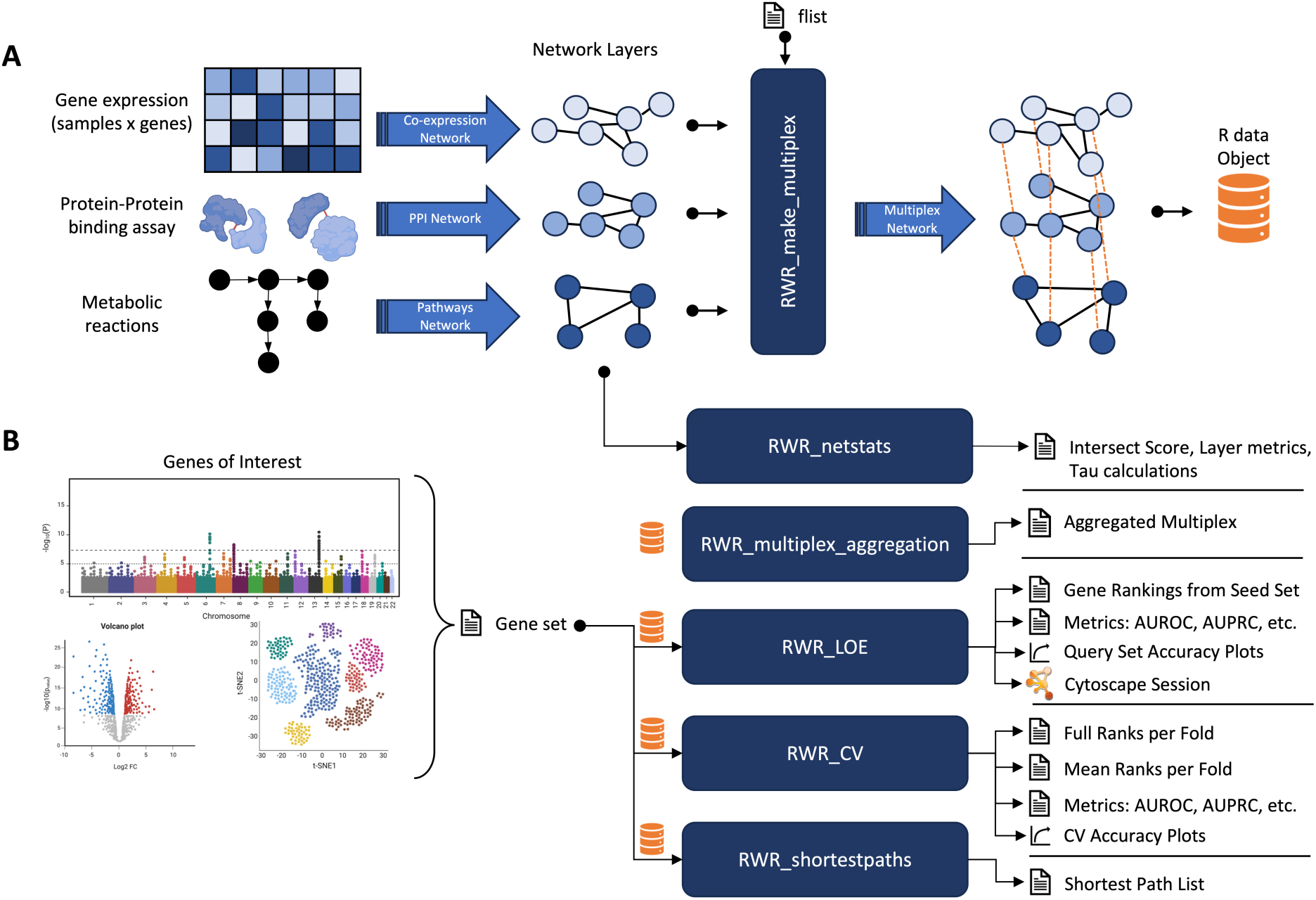
A general workflow for using the RWRtoolkit. **A.** Illustration of how a user can generate several network layers from different omics data sources, which become input to the RWRtoolkit workflow. Once the user has networks in the correct format, they can then refer to them via a flist file and use *RWR_make_multiplex* to turn them into a homogeneous multiplex network (e.g., multiple layers of gene-to-gene relationships). This multiplex is wrapped in an RData object that is saved for future use. **B.** A demonstration of how the user can now execute a variety of multi-omic analyses, most of which require the RData object as input. A set of genes of interest (gene set) from discovery studies such as GWAS or differential expression analysis can be used as input to multiple tools. These tools output a variety of files that show how functionally connected the genes in the gene set are to each other, or to a second gene set of interest, or to all the other genes in the multiplex. Some resulting networks can be automatically visualized in Cytoscape via the RCy3 R package (Gustavsen, 2019). Figure uses illustrations created with BioRender.com.

**Table 1.** RWRtoolkit Execution Commands. Examples of how to call each of the RWRtoolkit functions from either an R environment or a command line environment.

| Command | R Script | Shell Script |
| --- | --- | --- |
|  | Assumes RWRtoolkit has been installed and “example_data” has been set as working directory:<br>`library(RWRtoolkit) extdata.dir <- system.file(package="RWRtoolkit") setwd(extdata.dir)` | Relative paths assume the scripts are run from within the `inst` directory of the downloaded github repository or the main directory of the already installed R package directory |
| Make Multiplex | `RWR_make_multiplex(flist='./example_data/flist.tsv')` | `Rscript ./scripts/run_make_multiplex.R --flist ./example_data/flist.tsv` |
| Netstats | `RWR_netstats( data = './example_data/string_interactions.Rdata', outdir = './netstats_output', network_1 = './example_data/netstat/combined_score-random-gold.tsv', network_2 = './example_data/netstat/combined_score-random-test.tsv', basic_statistics = T, scoring_metric = "both", pairwise_between_mpo_layer = T, multiplex_layers_to_refnet = T, net_to_net_similarity = T, calculate_tau_for_mpo = T, calculate_exclusivity_for_mpo = T )` | `Rscript ./scripts/run_netstats.R \ --data ./example_data/string_interactions.Rdata \ --outdir ./netstats_output \ --network_1 ./example_data/netstat/combined_score-random-gold.tsv \ --network_2 ./example_data/netstat/combined_score-random-test.tsv \ --basic_statistics \ --pairwise_between_mpo_layer \ --multiplex_layers_to_refnet \ --net_to_net_similarity \ --calculate_tau_for_mpo \ --calculate_exclusivity_for_mpo --verbose` |
| Network Aggregation | `RWR_network_aggregation( data = './example_data/string_interactions.Rdata', outdir = './netstats_networks', merged_with_all_edges = T, merged_with_edgecounts = T )` | `Rscript ./scripts/run_network_aggregation.R \ --data ./example_data/string_interactions.Rdata \ --outdir ./netstats_networks \ --merged_with_all_edges \ --merged_with_edgecounts` |
| LOE | `RWR_LOE( data = './example_data/string_interactions.Rdata', seed_geneset = './example_data/geneset1.tsv', outdir = './loe_output_dir' )` | `Rscript ./scripts/run_loe.R \ --data ./example_data/string_interactions.Rdata \ --seed_geneset ./example_data/geneset1.tsv \ --outdir ./loe_output_dir` |
| CV | `RWR_CV( data = './example_data/string_interactions.Rdata', geneset_path= './example_data/geneset1.tsv', method='kfold', folds=3, outdir = './cv_kfold_output_dir' )` | `Rscript ./scripts/run_cv.R \ --data ./example_data/string_interactions.Rdata \ --geneset ./example_data/geneset1.tsv \ --method kfold \ --folds 3 \ --outdir ./cv_kfold_output_dir` |
| Shortest Paths | `RWR_ShortestPaths( data = './example_data/string_interactions.Rdata', source_geneset= './example_data/geneset1.tsv', target_geneset='./example_data/geneset2.tsv', outdir='./shortest_paths_output' )` | `Rscript ./scripts/run_shortestpaths.R \ --data ./example_data/string_interactions.Rdata \ --source_geneset ./example_data/geneset1.tsv \ --target_geneset ./example_data/geneset2.tsv \ --outdir ./shortest_paths_output` |

### 2.1 Multiplex Generation

RWRtoolkit usage typically starts with the *RWR_make_multiplex* command, which handles the creation of the multiplex network. It requires a descriptor file (known as an “flist”) that lists the full path to each network layer to be included in the multiplex. Each network layer’s file must be formatted as a simple delimited edge list with a column for the source genes, a column for the target genes, and an optional weight column (see **Supplementary Tables 1, 2, and 3** for file examples). The generated multiplex network is automatically saved as an Rdata object containing the individual layers as igraph [16,17] networks, the multiplex transition matrices, and network metadata. This Rdata object is used as input for most downstream commands.

### 2.2 Multiplex RWR Applications

#### 2.2.1 Network Layer and Multiplex Statistics

The contents of a multiplex network affect the outcomes of RWR analyses. The *RWR_netstats* command lets the user evaluate individual network layers or the contents of an entire multiplex network containing many layers, using a variety of options for calculating network metrics as well as functions for multiplex network evaluation. Basic statistics (basic_statistics) for individual layers can be calculated, as well as more complex relationships such as jaccard or overlap scores for inter-layer similarities (pairwise_between_mpo_layer), multiplex layer to reference network (multiplex_layers_to_refnet), and single network to network similarities (net_to_net_similarity). During a random walk, the tau parameter affects the probability of the walker visiting each specific layer, allowing the user to bias the walk to certain layers of higher importance. Users can supply their own tau, or use the calculate_tau function to return a tau value for each layer.

#### 2.2.2 Evaluating multiplex networks and gene sets using cross-validation

The predictive ability of a multiplex network can be determined using cross validation of gold standard or reference gene sets with the *RWR_CV* command. A gold standard gene set typically contains genes that are known to be functionally related (e.g., all are members of one biosynthetic pathway, all are annotated with the same GO/KEGG term, etc.). The hypothesis is that gold standard genes purposely left out from the seed set should be found with relatively high precision (i.e., highly ranked by RWR) if the underlying networks are indeed functionally predictive. The *RWR_CV* command allows the user to provide a gold standard gene set and use k-fold, leave-one-out, or singleton cross validation to score the ability to find the left out gene(s).

*RWR_CV* produces output files containing the RWR score and rank of each gene in the multiplex (as described by *RWR_LOE,* for each fold), mean rank of each gene across all folds, evaluation metrics based on the ranks of the left-out genes for each fold, and an evaluation summary file. Evaluation metrics, such as AUPRC and AUROC, can output as plots as well. File descriptions and examples can be found in Supplemental Tables 13-16.

#### 2.2.3 Ranking Genes Using Multiple Lines of Evidence (LOE)

The *RWR_LOE* command uses RWR to rank all genes in the multiplex network with respect to a gene-set of interest (seed genes), which provides multi-omic biological context for the seeds. The ranks and scores of all genes can be output to a file. A second gene set can be provided in order to evaluate the topological relationship between two sets of genes. When a second gene set is provided, those genes are flagged within the ranked output. The network context around the top ranked genes can be easily visualized via an integrated connection to Cytoscape [18] using the RCy3 [19] R package.

#### 2.2.4 Extracting The Shortest Paths Between Genes

In a network there can exist many unique paths between two particular nodes. Obtaining the shortest paths between any two given nodes within a network can provide crucial insight to a network’s topology or the relationship between those nodes. *RWR_ShortestPaths* calculates the pairwise shortest paths between source and target gene sets, and returns them as a series of edges that form the shortest path, the layers in which those edges exist, edge weights, and normalized edge weights.

#### 2.2.5 Network Aggregation Functions

Two methods of network aggregation are provided to merge the layers of a multiplex network into a single monoplex network. The *merged_with_all_layers* function aggregates all layers maintaining multiple edges between nodes. The *merged_with_all_edgecounts* function aggregates all layers of the multiplex, but instead edge weight is calculated as the sum of all shared edges within the multiplex network.

## 3. Results

We used RWRtoolkit to explore genetic relationships in two promising biofuel crops: switchgrass (*Panicum virgatum*) and pennycress (*Thlaspi arvense*). Since large scale multi-omic pennycress and switchgrass data are not yet available, we generated a multiplex network from varied data layers available for *Arabidopsis thaliana*, a model species which is in the same family as pennycress (**Table 2**).

**Table 2.**
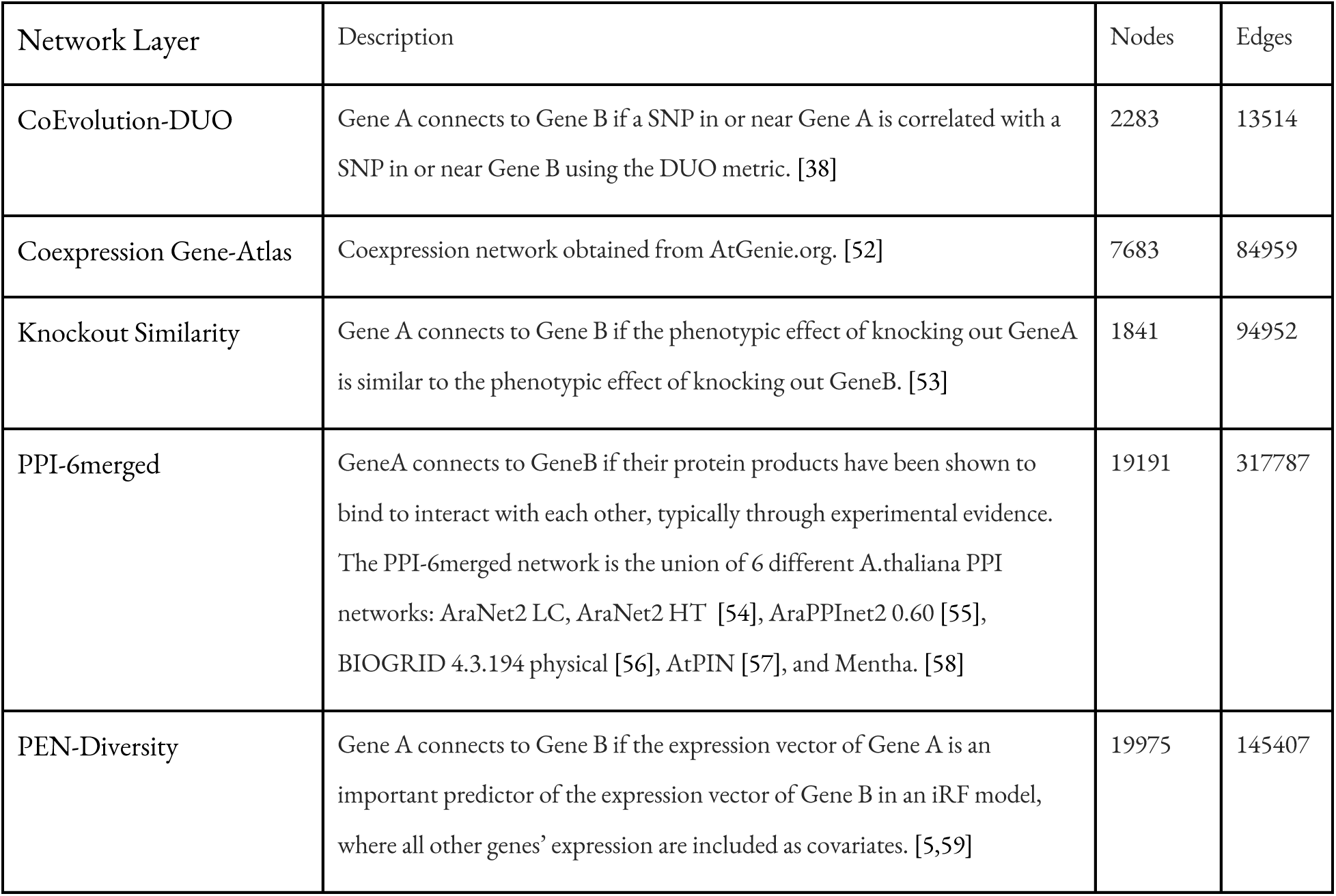
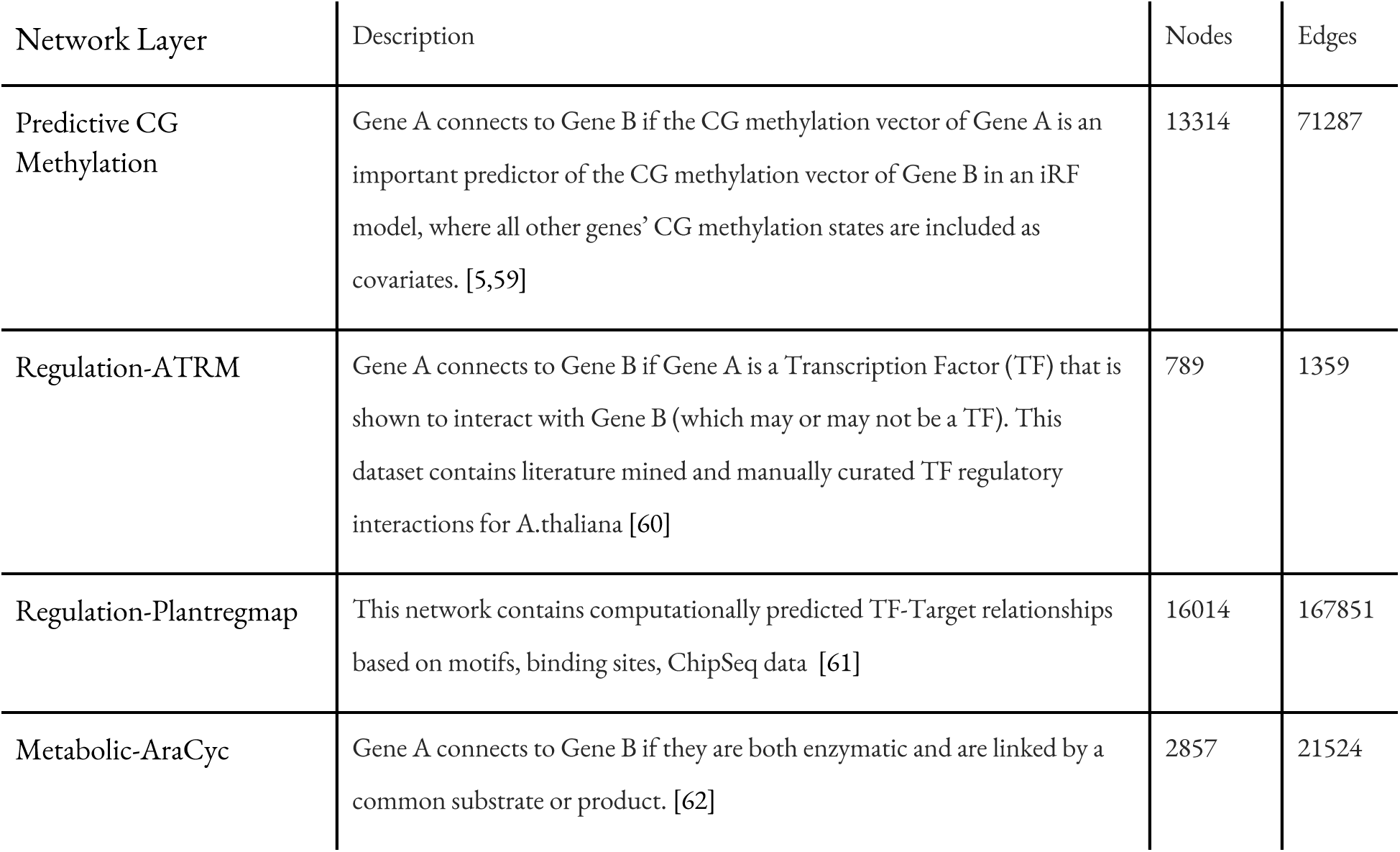
Multiplex Network Layers. A list of all network layers within the Comprehensive Multiplex Network.

### 3.1 Multiplex network validation using RWR and cross validation (*RWR_CV*)

We ran *RWR_CV* using k-fold (k=5) on each of 25 MAPMAN-derived [20] gene sets to validate the predictive ability of the multiplex network. This resulted in an average AUROC of 0.91 across all gene sets and folds, indicating a strong overall ability to find the left-out genes from a functional group and rank them highly. Conversely, when we performed the same analysis on 1000 randomly rewired multiplexes using command-line *RWR_CV*, the overall average AUROC was 0.49 (where AUROC of 0.50 is considered the equivalent of random). A comparison of AUROC densities for an individual MAPMAN gene set and all sets in total are illustrated in **figures 3a and 3b** (respectively). Individual *RWR_CV* comparison statistics for each MAPMAN gene set can be found in **Table 3**.

**Figure 3.**
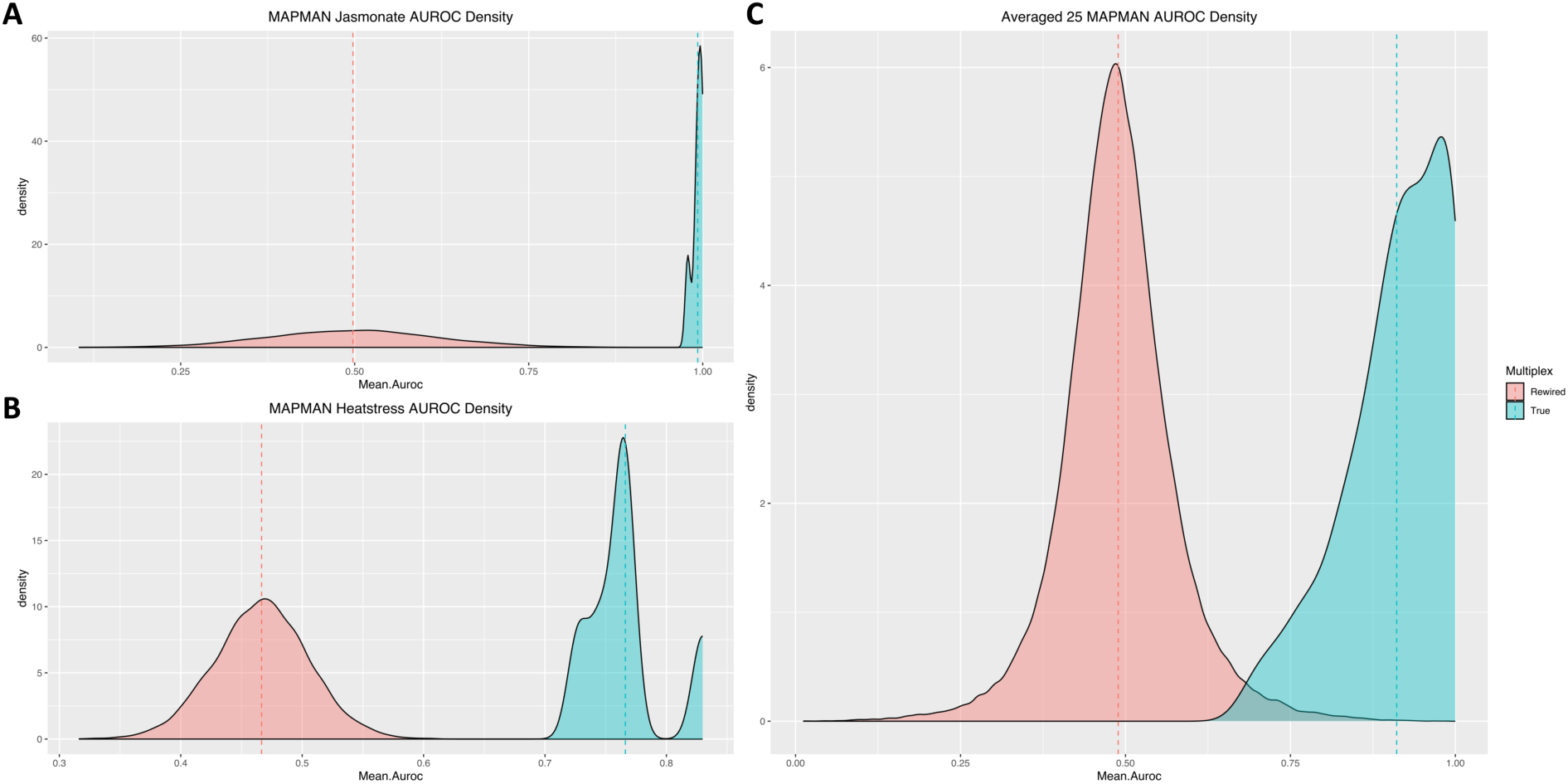
A comparison of the mean AUROC scores from the kfold output of *RWR_CV* using the true Comprehensive Multiplex (blue) and 1000 randomly rewired multiplex networks (red). **A.** Illustration of a comparison of AUROC density across 5 folds using an individual set of genes curated for Jasmonate signaling obtained from MAPMAN. The true comprehensive multiplex (blue) has a mean AUROC across 5 folds of 0.993 whereas the 1000 rewired multiplexes have an average mean AUROC across 5 folds of 0.498. **B.** Depiction of a comparison of AUROC density across 5 folds using an individual set of genes curated for Heat Stress signaling obtained from MAPMAN. The True comprehensive multiplex has a mean AUROC across 5 folds of 0.766. The average mean AUROC across 5 folds for the 1000 rewired multiplexes is 0.467. **C.** Illustration of a comparison of the average AUROC density across 25 gold standard gene sets generated from shared MAPMAN terms including Jasmonate and Heat Stress signaling. The true Comprehensive Multiplex has an overall average mean AUROC of 0.91 across all 25 gold standard gene sets, whereas the 1000 rewired multiplex networks have an overall average AUROC of 0.489 across all 25 gold standard gene sets, illustrating that the true Comprehensive Multiplex has meaningful biological connections compared to the completely random connections found across the 1000 rewired multiplex networks.

**Table 3.** Multiplex AUROC Comparison. This table contains the mean values and standard deviation of AUROC for *RWR_CV* Kfold cross validation for 25 curated gene sets which all share the same MAPMAN term. The average and standard deviation values for the rewired networks are averages and standard deviations across 1000 iterations of the *RWR_CV* Kfold Cross Validation in which the edges of each network were rewired for each iteration. Conversely, the average and standard deviation values for the Comprehensive network alone are those pertaining only to the average values across the single *RWR_CV* Kfold Cross Validation (k=5).

| Geneset | Rewired 1000 Average AUROC | Rewired 1000 std AUROC | Comprehensive net average AUROC | Comprehensive net std AUROC |
| --- | --- | --- | --- | --- |
| Absciscic acid | 4.989E-01 | 7.728E-02 | 8.710E-01 | 6.954E-02 |
| Auxin | 4.809E-01 | 4.295E-02 | 8.416E-01 | 3.975E-02 |
| Brassinosteroids | 4.891E-01 | 9.982E-02 | 9.393E-01 | 4.638E-02 |
| Cell Wall Synthesis | 5.008E-01 | 5.545E-02 | 9.815E-01 | 1.271E-02 |
| CHO metabolism | 4.968E-01 | 4.084E-02 | 9.388E-01 | 3.067E-02 |
| Coldstress | 5.008E-01 | 1.751E-01 | 9.812E-01 | 1.868E-02 |
| Cytokinin | 4.949E-01 | 1.269E-01 | 9.055E-01 | 7.550E-02 |
| Drought salt | 4.984E-01 | 8.276E-02 | 8.687E-01 | 3.908E-02 |
| Ethylene | 4.797E-01 | 6.081E-02 | 8.773E-01 | 4.811E-02 |
| Ethylene and EREBP | 4.929E-01 | 4.050E-02 | 9.248E-01 | 7.183E-03 |
| Fatty Acid | 4.912E-01 | 5.462E-02 | 9.318E-01 | 4.245E-02 |
| Flavonoids | 5.018E-01 | 7.089E-02 | 9.478E-01 | 2.879E-02 |
| Gibberelin | 4.776E-01 | 9.544E-02 | 8.496E-01 | 9.136E-02 |
| Glucosinolates | 4.987E-01 | 7.306E-02 | 9.394E-01 | 3.609E-02 |
| Heat stress | 4.665E-01 | 3.903E-02 | 7.662E-01 | 3.477E-02 |
| HeatshockTFs | 4.657E-01 | 3.859E-02 | 7.688E-01 | 5.851E-02 |
| Isoprenoids | 4.990E-01 | 5.616E-02 | 9.847E-01 | 1.511E-02 |
| Jasmonate | 4.975E-01 | 1.207E-01 | 9.928E-01 | 7.937E-03 |
| Lignin Biosynthesis | 5.026E-01 | 1.099E-01 | 9.990E-01 | 3.110E-04 |
| Major CHO | 4.899E-01 | 5.873E-02 | 9.135E-01 | 4.158E-02 |
| Phenylpropanoids | 5.007E-01 | 7.702E-02 | 9.970E-01 | 2.329E-03 |
| PS Light Reaction | 4.528E-01 | 4.627E-02 | 8.414E-01 | 3.272E-02 |
| PS Light Reaction II | 4.582E-01 | 7.547E-02 | 9.024E-01 | 1.358E-02 |
| Salicylic Acid | 5.006E-01 | 1.498E-01 | 8.992E-01 | 1.078E-01 |

### 3.2 Identifying Genes Contributing to switchgrass Well-Watered Shoot Biomass Using RWRtoolkit and KBase

Identifying genetic variants, genes, and biological pathways controlling plant biomass can prioritize gene targets for improving biofuel feedstocks. To this end, we performed a genome-wide association study (GWAS) for well-watered shoot biomass dry weight (**Table 4, Supplementary Fig 2**.) in the tetraploid bioenergy feedstock switchgrass (*Panicum virgatum*). Two GWAS models (BLINK and FarmCPU) identified 22 unique significant single nucleotide polymorphisms (SNPs) associated with well-watered shoot biomass at an FDR < 0.2. These were mapped to a total of 38 unique switchgrass genes based on genomic proximity.

**Table 4.** Switchgrass GWAS Hits. All significant loci found from the well watered dry weight shoot biomass GWAS.

| GWAS Methods | Chr Orig | Pos | Pvalue | MAF | nobs | FDR_Pvalue | Effect | DISTANCE | locusName | Phytozome Arabidopsis ortholog |
| --- | --- | --- | --- | --- | --- | --- | --- | --- | --- | --- |
| BLINK | 02K | 62843430 | 1.19E-13 | 2.30E-01 | 259 | 5.32E-07 | 4.77E-01 | 9199 | Pavir.2KG520800 | AT4G33925.1 |
| BLINK | 02K | 62843430 | 1.19E-13 | 2.30E-01 | 259 | 5.32E-07 | 4.77E-01 | 6514 | Pavir.2KG520900 | AT1G24625.1 |
| BLINK | 02K | 43638064 | 4.59E-13 | 1.93E-01 | 259 | 1.02E-06 | -4.77E-01 | -20510 | Pavir.2KG303000 | AT5G16560.1 |
| BLINK | 02K | 43638064 | 4.59E-13 | 1.93E-01 | 259 | 1.02E-06 | -4.77E-01 | -32665 | Pavir.2KG303302 | AT5G16550.1 |
| BLINK | 01N | 58593942 | 2.53E-12 | 1.68E-01 | 259 | 3.75E-06 | 5.04E-01 | -2241 | Pavir.1NG459600 | AT1G32450.1 |
| BLINK | 01N | 58593942 | 2.53E-12 | 1.68E-01 | 259 | 3.75E-06 | 5.04E-01 | 5266 | Pavir.1NG459619 |  |
| BLINK | 08N | 12302907 | 7.48E-12 | 5.02E-02 | 259 | 8.34E-06 | -7.39E-01 | -1715 | Pavir.8NG064200 | AT3G47570.1 |
| BLINK | 08N | 12302907 | 7.48E-12 | 5.02E-02 | 259 | 8.34E-06 | -7.39E-01 | 2805 | Pavir.8NG064201 |  |
| BLINK | 02K | 40684207 | 1.12E-11 | 1.89E-01 | 259 | 1.00E-05 | -5.13E-01 | -6514 | Pavir.2KG285100 | AT2G46210.1 |
| BLINK | 02K | 40684207 | 1.12E-11 | 1.89E-01 | 259 | 1.00E-05 | -5.13E-01 | 0 | Pavir.2KG286800 | AT2G46210.1 |
| BLINK | 01K | 46049518 | 2.84E-10 | 7.14E-02 | 259 | 2.11E-04 | -6.48E-01 | -12057 | Pavir.1KG445700 | AT1G12640.1 |
| BLINK | 01K | 46049518 | 2.84E-10 | 7.14E-02 | 259 | 2.11E-04 | -6.48E-01 | -12057 | Pavir.1KG445700 | AT1G12640.1 |
| BLINK | 01K | 46049518 | 2.84E-10 | 7.14E-02 | 259 | 2.11E-04 | -6.48E-01 | -12057 | Pavir.1KG445700 | AT1G12640.1 |
| BLINK | 01K | 46049518 | 2.84E-10 | 7.14E-02 | 259 | 2.11E-04 | -6.48E-01 | 12139 | Pavir.1KG445800 | AT5G19700.1 |
| BLINK | 06N | 36174982 | 1.19E-09 | 2.07E-01 | 259 | 7.60E-04 | -3.82E-01 | -4398 | Pavir.6NG117300 | AT1G60140.1 |
| BLINK | 06N | 36174982 | 1.19E-09 | 2.07E-01 | 259 | 7.60E-04 | -3.82E-01 | -6348 | Pavir.6NG117506 | AT1G23890.2 |
| FarmCPU | 02K | 40684207 | 1.88E-19 | 1.89E-01 | 259 | 8.38E-13 | -5.33E-01 | -6514 | Pavir.2KG285100 | AT2G46210.1 |
| FarmCPU | 02K | 40684207 | 1.88E-19 | 1.89E-01 | 259 | 8.38E-13 | -5.33E-01 | 0 | Pavir.2KG286800 | AT2G46210.1 |
| FarmCPU | 08N | 12302907 | 5.39E-16 | 5.02E-02 | 259 | 1.20E-09 | -6.31E-01 | -1715 | Pavir.8NG064200 | AT3G47570.1 |
| FarmCPU | 08N | 12302907 | 5.39E-16 | 5.02E-02 | 259 | 1.20E-09 | -6.31E-01 | 2805 | Pavir.8NG064201 |  |
| FarmCPU | 08K | 12795029 | 6.62E-14 | 1.22E-01 | 259 | 9.83E-08 | 4.36E-01 | -5583 | Pavir.8KG108500 | AT2G37990.1 |
| FarmCPU | 08K | 12795029 | 6.62E-14 | 1.22E-01 | 259 | 9.83E-08 | 4.36E-01 | 8491 | Pavir.8KG136900 | AT1G52190.1 |
| FarmCPU | 08K | 12795029 | 6.62E-14 | 1.22E-01 | 259 | 9.83E-08 | 4.36E-01 | 8491 | Pavir.8KG136900 | AT1G52190.1 |
| FarmCPU | 05K | 47363351 | 4.04E-13 | 6.18E-02 | 259 | 4.51E-07 | -5.89E-01 | -4419 | Pavir.5KG269007 |  |
| FarmCPU | 05K | 47363351 | 4.04E-13 | 6.18E-02 | 259 | 4.51E-07 | -5.89E-01 | 8439 | Pavir.5KG523200 |  |
| FarmCPU | 03K | 1996767 | 1.04E-10 | 1.68E-01 | 259 | 9.30E-05 | -3.02E-01 | 3444 | Pavir.3KG025900 |  |
| FarmCPU | 03K | 1996767 | 1.04E-10 | 1.68E-01 | 259 | 9.30E-05 | -3.02E-01 | -1626 | Pavir.3KG041300 | AT3G44735.2 |
| FarmCPU | 05N | 51191444 | 2.12E-10 | 1.20E-01 | 259 | 1.58E-04 | 4.32E-01 | 13229 | Pavir.5NG411500 | AT2G33250.1 |
| FarmCPU | 05N | 51191444 | 2.12E-10 | 1.20E-01 | 259 | 1.58E-04 | 4.32E-01 | 13229 | Pavir.5NG411500 | AT2G33250.1 |
| FarmCPU | 05N | 51191444 | 2.12E-10 | 1.20E-01 | 259 | 1.58E-04 | 4.32E-01 | -14760 | Pavir.5NG430240 |  |
| FarmCPU | 06K | 14705578 | 9.72E-10 | 1.91E-01 | 259 | 6.19E-04 | 2.76E-01 | -5601 | Pavir.6KG187400 | AT1G16840.4 |
| FarmCPU | 06K | 14705578 | 9.72E-10 | 1.91E-01 | 259 | 6.19E-04 | 2.76E-01 | 0 | Pavir.6KG188500 | AT1G16860.1 |
| FarmCPU | 06K | 14705578 | 9.72E-10 | 1.91E-01 | 259 | 6.19E-04 | 2.76E-01 | 0 | Pavir.6KG188500 | AT1G16860.1 |
| FarmCPU | 06K | 14705578 | 9.72E-10 | 1.91E-01 | 259 | 6.19E-04 | 2.76E-01 | 0 | Pavir.6KG188500 | AT1G16860.1 |
| FarmCPU | 06K | 14705578 | 9.72E-10 | 1.91E-01 | 259 | 6.19E-04 | 2.76E-01 | 0 | Pavir.6KG188500 | AT1G16860.1 |
| FarmCPU | 09K | 52780266 | 4.18E-09 | 1.85E-01 | 259 | 2.33E-03 | 6.12E-01 | -9455 | Pavir.9KG406700 | AT4G17910.1 |
| FarmCPU | 09K | 52780266 | 4.18E-09 | 1.85E-01 | 259 | 2.33E-03 | 6.12E-01 | -4933 | Pavir.9KG406800 | AT2G02970.1 |
| FarmCPU | 09K | 52780266 | 4.18E-09 | 1.85E-01 | 259 | 2.33E-03 | 6.12E-01 | -4933 | Pavir.9KG406800 | AT2G02970.1 |
| FarmCPU | 01K | 21997091 | 1.21E-08 | 3.44E-01 | 259 | 5.99E-03 | 2.92E-01 | 9147 | Pavir.1KG276300 | AT5G43820.1 |
| FarmCPU | 01K | 21997091 | 1.21E-08 | 3.44E-01 | 259 | 5.99E-03 | 2.92E-01 | 9147 | Pavir.1KG276300 | AT5G43820.1 |
| FarmCPU | 01K | 21997091 | 1.21E-08 | 3.44E-01 | 259 | 5.99E-03 | 2.92E-01 | 9147 | Pavir.1KG276300 | AT5G43820.1 |
| FarmCPU | 01K | 21997091 | 1.21E-08 | 3.44E-01 | 259 | 5.99E-03 | 2.92E-01 | 13972 | Pavir.1KG277500 | AT5G24500.1 |
| FarmCPU | 01K | 21997091 | 1.21E-08 | 3.44E-01 | 259 | 5.99E-03 | 2.92E-01 | 13972 | Pavir.1KG277500 | AT5G24500.1 |
| FarmCPU | 01K | 21997091 | 1.21E-08 | 3.44E-01 | 259 | 5.99E-03 | 2.92E-01 | 13972 | Pavir.1KG277500 | AT5G24500.1 |
| FarmCPU | 05K | 60634281 | 1.35E-08 | 5.79E-02 | 259 | 6.02E-03 | -3.57E-01 | -15923 | Pavir.5KG758000 | AT2G41510.1 |
| FarmCPU | 05K | 60634281 | 1.35E-08 | 5.79E-02 | 259 | 6.02E-03 | -3.57E-01 | -1975 | Pavir.5KG758200 | AT1G50460.1 |
| FarmCPU | 02N | 35440741 | 1.66E-08 | 1.20E-01 | 259 | 6.73E-03 | 2.84E-01 | -4118 | Pavir.2NG156300 | AT4G02780.1 |
| FarmCPU | 02N | 35440741 | 1.66E-08 | 1.20E-01 | 259 | 6.73E-03 | 2.84E-01 | 16670 | Pavir.2NG156400 | AT4G02780.1 |
| FarmCPU | 06K | 39449007 | 2.35E-08 | 4.73E-01 | 259 | 8.71E-03 | -1.86E-01 | 0 | Pavir.6KG295706 |  |
| FarmCPU | 06K | 39449007 | 2.35E-08 | 4.73E-01 | 259 | 8.71E-03 | -1.86E-01 | 360 | Pavir.6KG295800 | AT3G12050.1 |
| FarmCPU | 01K | 39637992 | 7.01E-08 | 3.92E-01 | 259 | 2.40E-02 | -2.19E-01 | 0 | Pavir.1KG366900 | AT1G25580.1 |
| FarmCPU | 01K | 39637992 | 7.01E-08 | 3.92E-01 | 259 | 2.40E-02 | -2.19E-01 | 3678 | Pavir.1KG367100 | AT3G27540.1 |
| FarmCPU | 02N | 6201763 | 1.62E-07 | 8.88E-02 | 259 | 5.17E-02 | -2.90E-01 | 24 | Pavir.2NG060000 | AT3G09630.1 |
| FarmCPU | 02N | 6201763 | 1.62E-07 | 8.88E-02 | 259 | 5.17E-02 | -2.90E-01 | 3414 | Pavir.2NG060546 | AT3G14470.1 |
| FarmCPU | 05K | 15910143 | 1.99E-07 | 9.07E-02 | 259 | 5.92E-02 | -3.40E-01 | -4196 | Pavir.5KG216014 |  |
| FarmCPU | 05K | 15910143 | 1.99E-07 | 9.07E-02 | 259 | 5.92E-02 | -3.40E-01 | -4196 | Pavir.5KG216014 |  |
| FarmCPU | 05K | 15910143 | 1.99E-07 | 9.07E-02 | 259 | 5.92E-02 | -3.40E-01 | 0 | Pavir.5KG216021 | AT3G62160.1 |
| FarmCPU | 09N | 46512678 | 2.60E-07 | 1.91E-01 | 259 | 7.25E-02 | 2.14E-01 | 12848 | Pavir.9NG277800 |  |
| FarmCPU | 09N | 46512678 | 2.60E-07 | 1.91E-01 | 259 | 7.25E-02 | 2.14E-01 | 11217 | Pavir.9NG413642 |  |
| FarmCPU | 09K | 6063075 | 6.81E-07 | 3.11E-01 | 259 | 1.79E-01 | 3.16E-01 | -653 | Pavir.9KG224800 | AT2G45490.1 |
| FarmCPU | 09K | 6063075 | 6.81E-07 | 3.11E-01 | 259 | 1.79E-01 | 3.16E-01 | -5059 | Pavir.9KG224900 | AT1G47310.1 |

We next wanted to understand the biological context among our Switchgrass well-watered shoot biomass gene set. However, there are limited publicly available networks that describe gene-gene relationships in switchgrass [21,22], so we mapped the 38 switchgrass GWAS genes to 32 Arabidopsis (*Arabidopsis thaliana*) orthologs using Phytozome [23]. For a greater understanding of the functional context of these GWAS results we leveraged our Arabidopsis multiplex network consisting of nine distinct lines of biological evidence to explore the relationships among these orthologs (**Fig. 4A**).

**Figure 4.**
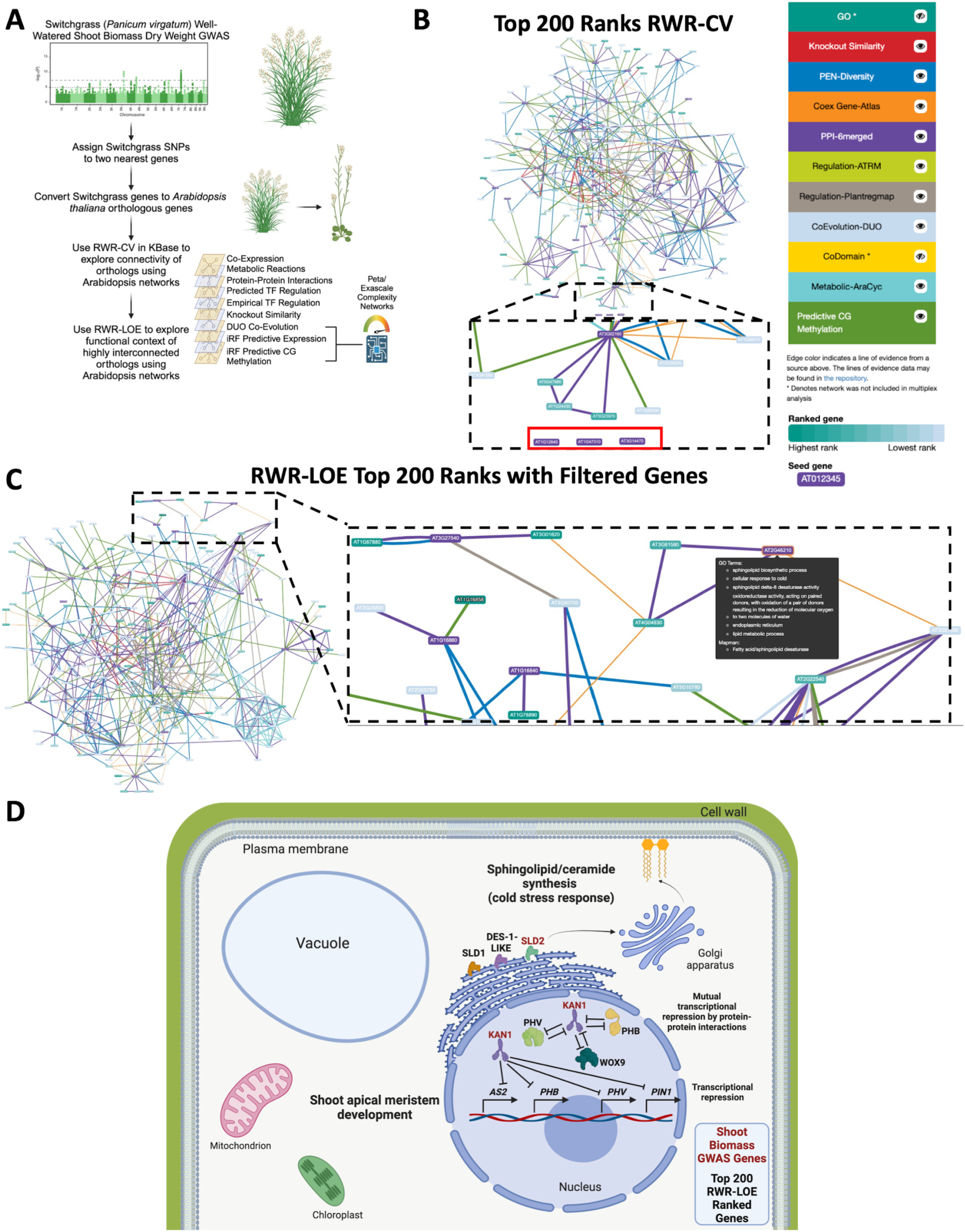
Exploring switchgrass GWAS results using Arabidopsis multiplex networks in KBase using RWRtoolkit. **A.** Workflow diagram. Switchgrass GWAS models identify significant single nucleotide polymorphism (SNP) associations with well-watered shoot biomass dry weight, and these SNPs are assigned to the two nearest Switchgrass genes. Switchgrass genes are converted to Arabidopsis orthologs, which are used as seeds in Arabidopsis multiplex networks. KBase apps were used to explore highly interconnected genes and identify functional context of the GWAS genes using a 9-layer multiplex, including 3 network layers (DUO Co-Evolution, iRF Predictive Expression, and iRF Predictive CG Methylation) derived from high-performance computing using models with petascale or exascale-level combinatorial complexity. Figure created using Biorender.com. **B.** Using KBase, *RWR-CV* identified three seed genes (inset, red box) that were unconnected from the top 200 ranked genes. Seed genes (Arabidopsis orthologs from GWAS results) are displayed as purple nodes, and teal to light blue nodes are color scaled based on *RWR-LOE* ranks. Edge colors indicate the line of evidence from which each gene-gene relationship was derived. **C.** Visualization of KBase *RWR-LOE* top 200 ranks output using orthologs of genes that were highly interconnected based on *RWR-CV* (“filtered genes”). Inset: Magnified view of network. When hovering over a gene, users can view Gene Ontology (GO), knockout phenotype and MAPMAN annotations for that gene. **D.** Framework for a conceptual model of Switchgrass well-watered shoot biomass with GWAS genes (red) and top 200 ranked genes by *RWR-LOE* (black). Genes from GWAS and *RWR-LOE* ranks implicated shoot apical meristem development, homeodomain transcription factors involved in transcriptional repression, and sphingolipid/ceramide synthesis. Figure made with BioRender.com.

*RWR_CV* and *RWR_LOE* functionality and the Arabidopsis multiplex network were incorporated into the DOE Systems Biology Knowledgebase (KBase [15]) interface (**Fig. 4B-C**). Using the visualization derived from the KBase tools, we first removed three orthologs (AT1G12640, AT1G47310, AT3G14470) that were not connected to any other GWAS genes, nor to the top 200 genes ranked by *RWR_CV* with 5-fold cross validation (**Fig. 4B**). We then used the remaining 29 GWAS gene orthologs as seeds for the *RWR_LOE* application to help generate a framework for a conceptual model depicting the top candidate genes from GWAS and their functional context derived from other high ranking genes (**Fig. 4D**). Here, we focus our discussion on two genes identified by shoot biomass GWAS and their relevant *RWR_LOE* connections.

Both BLINK and FarmCPU GWAS models identified the SNP Chr02K_40684207 as significant, with FDR-adjusted p-values of 1.00E-5 and 8.38E-13, respectively (**Table 4, Supplementary Fig. 2**). This SNP is located in the switchgrass gene Pavir.2KG286800, which is orthologous to the Arabidopsis gene *SPHINGOID LCB DESATURASE 2* (*SLD2*, AT2G46210.1) (**Table 4**). SLD2 plays a crucial role in sphingolipid biosynthesis by catalyzing the desaturation of long-chain bases (LCBs) at position 8 [24]. *RWR_LOE* analysis revealed interactions between *SLD2* and SPHINGOID LCB DESATURASE 1 (SLD1, AT3G61580) and DES-1-LIKE (AT4G04930), which are delta 8 and delta 4 desaturases through the PPI layer (**Fig. 4D**), as well as AGL18, a MADS-domain transcription factor (AT3G57390) through the coexpression layer.

Additionally, the BLINK GWAS model identified Chr02K_43638064 as significant (FDR-adjusted p-value 1.02E-06, **Table 4**). The nearest gene to this SNP is Pavir.2KG303000, which is orthologous to AT5G16560.1, or *KANADI1* (*KAN1)*. *KAN1* is a transcription factor (TF) with a significant role in adaxial-abaxial (top and bottom) polarity in leaves and the proper development of the shoot apical meristem (SAM) via auxin signaling through interactions with auxin-related genes [25]. Previous work implicated *KAN1* in a growth-defense tradeoff regime in *Arabidopsis thaliana* through jasmonic acid (JA) signaling [26]. The authors reported that JA activates *KAN1* which suppresses auxin biosynthesis, transport, and signaling, ultimately inhibiting growth [26]. Importantly, this suggests that SNPs affecting KAN1 may alter growth and biomass phenotypes. To gain a more comprehensive understanding of the mechanisms involved in this regulatory network, we examined the genes connected to *KAN1* in the KBase *RWR_LOE* Narrative.

Visualizations of the lines of evidence around *KAN1* and within multiplex network revealed protein-protein interactions (PPI-6merged, **Table 2**) between products of *KAN1* and *PHAVOLUTA* (*PHV*), *PHABULOSA* (*PHB*), and *WUSCHEL-RELATED HOMEOBOX 9* (*WOX9*; **Fig. 4D**) [27]. *KAN1* was also connected to *PHV*, *PHB*, ASYMMETRIC LEAVES 2 (*AS2*) and PIN-FORMED 1 (*PIN1*) through TF regulatory interactions (Regulation-ATRM, described in **Table 2**). Additionally, the results of *RWR_LOE* exhibited a machine learning-predicted epigenetic relationship (Predictive CG Methylation, **Table 2**) between *KAN1* and *WOX9.* Together, the output of *RWR_CV* and *RWR_LOE* identified shoot apical meristem development, long-chain fatty acid modifications, and homeodomain transcription factors as strong candidates affecting shoot biomass.

### 3.3 Predicting the Functional Effects of Gene Edits

Next, we applied RWRtoolkit to explore biological pathways surrounding two distinct genetically modified lines of pennycress (*Thlaspi arvense*). Pennycress is a cover crop in the Brassicaceae family with great potential to produce biodiesel and sustainable aviation fuel through large seed yields containing high volumes of long-chain fatty-acids. Jarvis et al. [13] recently demonstrated that pennycress lines with dual knockout of genes *FAE1* and *FAD2* produced seeds with 91% oleic acid content (significantly improved from 12% wild type accumulation), but at the cost of significantly reduced seed yield and stunted plant growth. Interestingly, the authors found that an alternative dual knockout of *FAE1* and *ROD1* accumulated up to 60% oleic acid with no obvious growth deficit. Here we used RWRtoolkit with an Arabidopsis multiplex network to investigate the difference in functional context between the *FAE1* and *FAD2* knockouts (*FAE1/FAD2*) and the *FAE1* and *ROD1* knockouts (*FAE1/ROD1*).

We ran *RWR_LOE* for each knockout pair separately and noted that many of the top 200 ranked genes from the *FAE1/ROD1* run remained very highly ranked in the *FAE1/FAD2* run. However, some of the top 200 genes fell drastically in ranking, indicating that they were no longer part of the same functional context after swapping *ROD1* for *FAD2* in the LOE runs (Illustrated in **Supplemental Figure 3**). Such “differentially ranked” genes can be considered as candidates driving functional changes that result in the phenotypic differences observed between the two dual knockout lines.

Using the set differential methodology to illustrate these differential rank differences, Gene Ontology enrichment of the intersection of the top 200 ranked genes from the *FAE1/FAD2* seeds and the *FAE1/ROD1* seeds (i.e., genes ranked highly by *RWR_LOE* for <u>both</u> knockout pairs) exhibited enrichment for fatty acid biosynthesis, fatty acid metabolic process, and sphingolipid metabolic process. Genes ranked highly by *RWR_LOE* for *FAE1/ROD1* but not for *FAE1/FAD2* showed three terms enriched: seed oil biogenesis, response to freezing, and lipid storage. The genes ranked highly by *RWR_LOE* for *FAE1/FAD2* but not for *FAE1/ROD1*, however, showed enrichment for multiple GO BP and KEGG terms beyond those expected for lipids, including **photoinhibition, regulation of circadian rhythm, and chloroplast rRNA processing**.

## 4. Discussion

Here, we demonstrate that RWRtoolkit enables the discovery of gene-to-gene relationships not previously apparent within a monoplex network topology, as well as gene-to-gene relationships across the broader multiplex surrounding a gene set of interest. Using the RWRtoolkit package, users can create and validate multiplex biological networks encoding multiple lines of evidence. Importantly, RWRtoolkit is agnostic to organism, tissue or condition. The user may explore biological pathways in non-model organisms by either using orthologs and available networks from model organisms as demonstrated in the present work by building custom networks from experimental data, or a combination of both approaches. In addition to obtaining topologically relevant gene-to-gene relationships from the multiplex networks, users can identify the lines of evidence driving these interactions (e.g., co-expression, protein-protein interactions, etc.) using Cytoscape or KBase. Additionally, peta/exascale-complexity networks derived from AI-based methods can be used as layers within the multiplex to identify relationships for poorly-annotated genes, including proteins of unknown function.

RWRtoolkit was designed as a user-friendly package for researchers familiar with R software and command line interfaces. Users who want to generate custom multiplex networks and use the entire suite of functions in RWRtoolkit can find the open-source code and vignettes on GitHub. For users who prefer a point-and-click graphical user interface or have limited bioinformatic experience, we have included *RWR_LOE* and *RWR_CV* as applications within KBase and provided pre-assembled *Arabidopsis thaliana* multiplex networks.

RWRtoolkit explores topological connectivity between seed genes and other genes based on multiple lines of evidence. In doing so, RWRtoolkit facilitates interpretation of a gene set outside of gene set enrichment analysis, with the goal of expanding the biological context of genes in a gene set which may not have been previously studied in the same experimental context. Moreover, the biological context between any group of genes is explainable based on the various types of biological evidence present in a multiplex network.

### 4.1 Multiplex Network Validation

First, we showed the predictive capacity of our Arabidopsis multiplex network by demonstrating that MAPMAN gene sets were highly interconnected when using network layers from data sources distinct from MAPMAN. By using ontology-derived, “gold standard” gene sets of interrelated genes from random walk exploration of a multiplex, we can ensure that our multiplex networks represent true biological connections, rather than random connections. Moreover, our Arabidopsis multiplex significantly outperformed recall of MAPMAN gene sets compared to permutations of randomly connected networks, indicating that this is a valid way to test whether a multiplex network contains biologically meaningful edges. We then demonstrated these applications in a real-world biological context using genesets derived from GWAS results and a gene editing experiment.

### 4.2 Well Watered Shoot Biomass GWAS Results

We applied the KBase *RWR_LOE* application to functionally contextualize GWAS results from the bioenergy feedstock switchgrass, as GWAS results for complex traits are often difficult to interpret because significant SNPs can map to a set of genes with largely uncharacterized relationships. *RWR_LOE* captured connections surrounding both sphingolipid production and a regulatory subnetwork of cell differentiation and specification that likely affects vascular development in the SAM. In addition, the interactions captured by RWR-LOE led to the development of a conceptual model framework highlighting these findings (**Fig. 4 D**).

Sphingolipids are integral to various cellular, developmental, and stress-related processes [28]. Though the SNP associated with SLD2 was identified by two GWAS models, SLD2 knockout experiments in *A. thaliana* showed no phenotype growth defects under normal conditions [24]. However, double mutants of *sld1 sld2* exhibited altered growth phenotypes under cold stress conditions accompanied by changes in the distribution of complex sphingolipids such as glucosyl-ceramide (GluCer) and GIPCs [24]. These findings suggest an ambiguous role for SLD2 in shoot biomass accumulation. However, AGL18, which was connected to SLD2 via coexpression, is essential in regulating the transition from vegetative to reproductive growth in plants [29] suggesting its involvement in shoot biomass development. Notably, the additional context provided by *RWR_LOE* highlights a connection between SLD2 and shoot biomass accumulation, a relationship that was not apparent from the GWAS results alone.

With respect to the SAM, the abaxial-adaxial regulatory network involved in shoot patterning and vascular development is primarily controlled by the *KANADI* TFs as well as the the Class III Homeodomain Leucine-Zipper (HD-ZIP III) TFs, *PHB* and *PHV* [30]. *KAN1* engages in both direct protein-protein interactions with *PHB* and *PHV* and influences their transcriptional activities antagonistically to preserve the required abaxial/adaxial boundary in apical meristem establishment and leaf development [31,32]. *AS2* is an important *LOB*-domain containing an adaxial regulator required for symmetrical leaf expansion [33]. *AS2* and *KAN1* are mutual transcriptional repressors controlling the lateral expansion and flatness of leaves which is fundamental to proper vegetative growth in the SAM [34]. Similarly, *WOX9* is a *WUS* homeobox-containing TF required for growth and maintenance of the vegetative SAM, in part through maintaining the population of undifferentiated stem cells [35]. *KAN1* engages in a protein-protein interaction with *WOX9*, likely to balance stem cell maintenance and cell fate/identity during vascular development in the SAM [27,36]. The methylation state of *KAN1* was found to be an important predictor of the methylation state of *WOX9*, suggesting an epigenetic relationship between these regulators in the SAM. *KAN1* also regulates key auxin-transport genes, such as *PIN1*, to orchestrate organ patterning and vascular development in the SAM. Specifically, *KAN1* directly inhibits *PIN1* by binding to a specific site downstream of *PIN1*, effectively restricting auxin flow by *PIN1* repression [31]. Additionally, the protein products of *WOX9*, *PHB*, and *PHV* were all shown to interact via a protein-protein interaction layer of the multiplex, further indicating a tightly interconnected regulatory network among these genes.

*RWR_LOE* significantly enhances traditional GWAS by uncovering additional topological connections within a multiplex (or monoplex) that would otherwise remain unknown. This approach enabled us to construct a conceptual model that includes a network of genetic influences on shoot biomass, which would not be possible with the GWAS results alone. We demonstrated how users can leverage the capabilities of *RWR_LOE* across a network to reveal mechanistic interactions and interpretations surrounding a user-defined gene set.

### 4.3 Exploring Dual Knockouts with *RWR_LOE*

We used RWRtoolkit with multi-omic Arabidopsis networks to understand the functional difference between *FAE1/ROD1* and *FAE1/FAD2* knockout plants from pennycress gene editing experiments. Using *RWR_LOE*, we aimed to explain the observed phenotypic differences between these genotypes based on differential network connectivity. The genes found in common in the top 200 ranks for both *FAE1/ROD1* and *FAE1/FAD2 RWR_LOE* runs exhibited enrichment for fatty acid biosynthesis-related GO and KEGG terms, which was expected given the well-described function of the three targeted genes in fatty acid biosynthesis and modification. The *FAE1/ROD1* specific subnetwork (i.e., genes not found in the *FAE1/FAD2* run) did not capture any obvious function beyond additional terms related to fatty acid synthesis and storage. However, the enriched GO and KEGG terms for the *FAE1/FAD2* subnetwork suggest impacts to the growth and development of the plant, and are possibly affected in a regulatory manner when *FAE1* and *FAD2* are knocked out, but not when *FAE1* and *ROD1* are knocked out. AT2G33800 (*EMB3113*), rank 160 in the *FAE1/FAD2* run, is annotated with the enriched GO BP term “chloroplast rRNA processing,” and has been shown to express a reduction in growth compared to wild type [37], offering a potential explanation as to why there exists growth reduction in the *FAE1/FAD2* knockout line.

To better understand the connectivity between *FAE1* and *FAD2* to *EMB3113*, we used *RWR_ShortestPaths* to explore the connections between these genes of interest. FAE1is connected to *EMB3113* through AT2G34315 via a GeneAtlas co-expression edge, and from AT2G34315 to *AT3G61920* via a DUO computed similarity (an advanced correlation metric) [38] edge. Finally, it connects *EMB3113* from *AT3G61920*, also via the DUO similarity metric (Fig 5 D). *FAD2* connects to *EMB3113* via a PEN (Predictive Expression Network) edge with *AT1G09750*.

**Figure 5.**
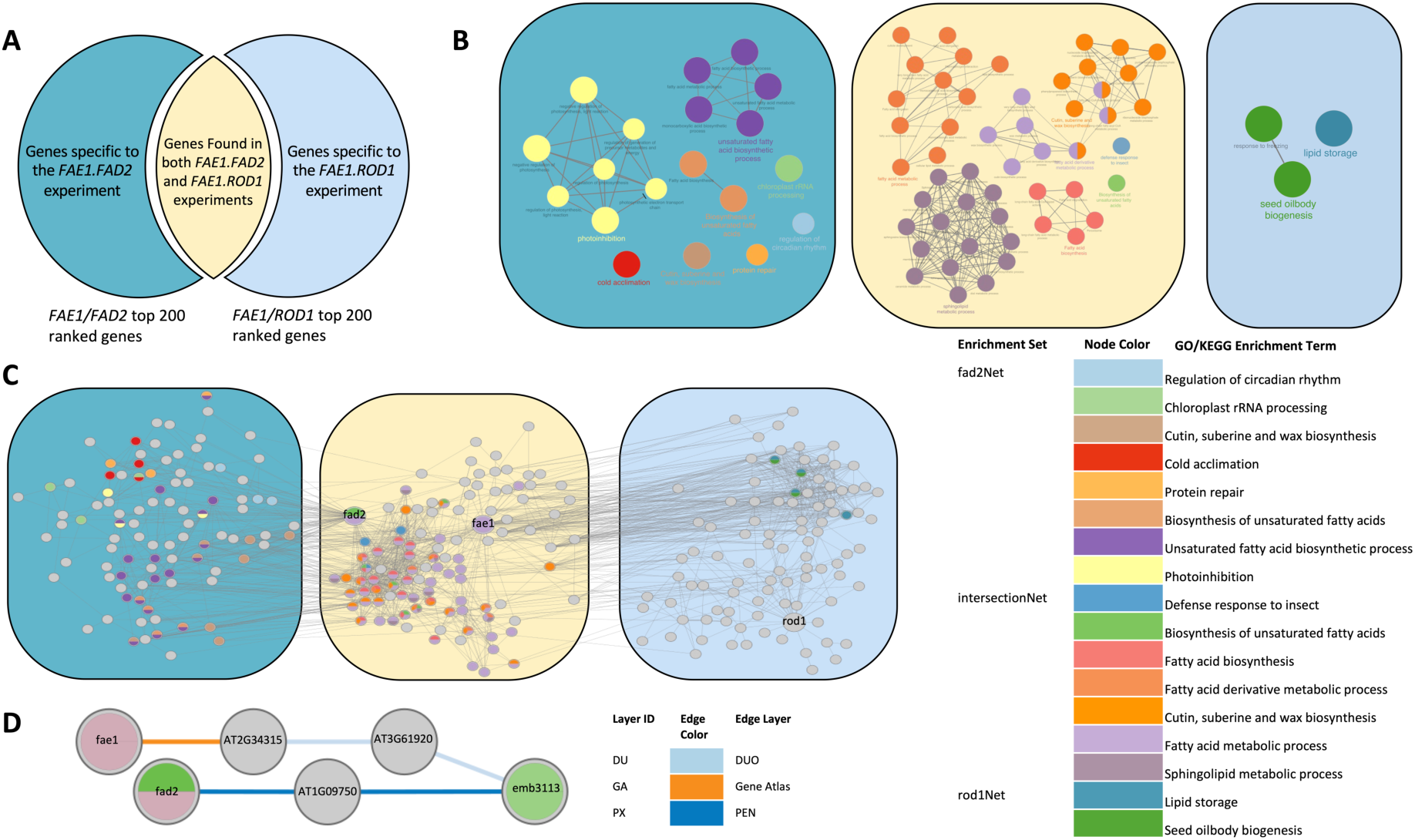
Top-ranked genes identify overlapping and distinct biological processes in pennycress from alternative knockout gene pairs. **A.** An illustration of the outcomes of using two knockout gene pairs (*FAE1/FAD2* and *FAE1/ROD1*) as seed genes for separate *RWR_LOE* runs. Genes under further investigation were either high ranking (top 200) for one knockout pair but not the other (blue), or high ranking for both (yellow) **B.** ClueGO enrichment of the three venn sets. Nodes are enrichment terms and edges represent term to term similarity defined by a corrected Cohen’s kappa statistic. **C.** The union of networks formed by the *FAE1.FAD2* top 200 genes and the *FAE1.ROD1* top 200 genes. From left to right: the FAD2 specific subnetwork, the intersection subnetwork (i.e., genes common to both *FAE1.FAD2* and *FAE1.ROD1*), and the ROD1 specific subnetwork. Genes (nodes) are colorized by their corresponding ClueGO enrichments. **D.** RWR Shortest paths between the *FAE1/FAD2* gene set and *EMB3113*. The shortest paths found from the *FAD2* to *EMB3113* uses edges from the predictive expression network while the path from *FAE1* to *EMB3113* uses edges from the gene atlas and duo network. In cases where edges exist on multiple layers, multiple labeled edges will exist between the two nodes.

Interestingly, AT2G34315 acts as an avirulence induced gene (*AIG1*)[39] and *AT3G61920* encodes a PADRE protein that exhibits downregulation when exposed to Pst DC3000 AvrRPS4 [40]. Therefore, genes within this path may suggest a connection between the growth/defense tradeoff in plant development. AT1G09750, the connecting gene between *FAD2* and *EMB3113*, encodes a metabolic enzyme with roles in hydrotropic response signal transduction and osmotic equilibrium maintenance [41]. Additionally, AT1G09750 has been quantified as being expressed during active growth and growth arrest developmental stages of Arabidopsis hypocotyls. Given AT1G09750’s role during active growth stages of development, we can begin to paint a more comprehensive picture as to why these pleiotropic effects occurred.

Using RWRtoolkit’s *RWR_LOE* and *RWR_ShortestPaths* functionality to explore the surrounding network topology of these knockout gene pairs, we start to gain some insights as to why the *FAE1/FAD2* dual knockout showed deleterious effects upon the growth of pennycress.

While the present manuscript focuses on gene-gene homogeneous networks, users could expand RWRtoolkit to include additional omic data such as metabolite-metabolite networks as well as heterogeneous networks (phenotype-gene, phenotype-metabolite, etc.). The focus of RWRtoolkit is intended to be applied to biological networks, but given that networks are domain agnostic and can signify any entity to entity relationship (including social networks, transportation networks, etc.), RWRtoolkit’s algorithms could be applied to any network data to identify highly ranked nodes using random walk with restart.

Together, we show that RWRtoolkit is an easy-to-use software package and KBase application that facilitates biological interpretation of experimental data sets using network analyses. We hope this package will provide another useful tool for researchers to interpret functional context from newly derived experimental data in order to accelerate scientific discovery.

## 5. Methods

### 5.1 Multiplex Construction and Layers

We constructed our comprehensive *Arabidopsis thaliana* multiplex consisting of 9 layers, defined in Table 2. All input networks were converted to be unweighted, and the multiplex constructed with a delta value of 0.5. The multiplex and its corresponding information can be found on GitHub (Comprehensive Multiplex, https://github.com/dkainer/RWRtoolkit-data/tree/main).

### 5.2. Network Validation

To ensure that our multiplex network was well constructed, we used *RWR_CV* with a series of well curated genes known to be highly connected within a biological system. These highly connected gene sets were derived from shared MAPMAN terms [20]. In order to ensure the validity of our multiplex network, we ran kfold cross validation (k=5) for each MAPMAN derived gene set. We compared the outputs of *RWR_CV* with another *RWR_CV* run using the same gene sets but on 1000 randomly rewired multiplexes with the same number of nodes and edges maintained in each layer.

### 5.3. GWAS

#### 5.3.1 SNP Variant Calling and Filtering

Variant calling methods for the SNPs were described previously [42]. Briefly, Illumina HiSeq X10 and Illumina NovaSeq 6000 paired-end sequencing at Department of Energy Joint Genome Institute and the HudsonAlpha Institute for Biotechnology were used for whole genome re-sequencing of the 260 *P. virgatum* genotypes. The median sequencing depth was 59x. The raw SNP dataset was filtered down to 4,458,778 SNPs for GWAS and all other downstream analyses: SNPs with more than 10% missing genotypes, genotypes with more than 10% missing SNPs, SNPs with severe departure from Hardy Weinberg Equilibrium (SNPs with HWE p-value < 1E-50), and SNPs with minor allele frequency < 0.05, SNPs with LD r2 >=0.07 were removed.

#### 5.3.2 Phenotyping

The aboveground shoot dry biomass was measured on 1442 *P. virgatum* plants (298 unique genotypes) grown under well-watered conditions in a greenhouse in multiple batches. Phenotypic outliers in the dataset were removed using the Median Absolute Deviation (MAD) method [43] with a MAD distance of 6 used as a threshold for removing outliers. Best Linear Unbiased Predictors (BLUPs) [44] of each genotype were obtained by running a linear model with genotypes as the random effect and the Batch as the fixed effect (covariate).

#### 5.3.3 GWAS

Association of the SNPs in the genome with the phenotypic trait (BLUPs) were calculated using GAPIT version 3 R package [45] with the following GWAS models: MLM [46], MLMM [47], FarmCPU [48], and BLINK [49]. The SNPs from the association test that passed the FDR threshold of 0.2 were considered significant. The significant SNPs were mapped to the two nearest *Panicum virgatum* genes, upstream and downstream using Version 5.1 snpEff annotation [15].

### 5.4 : Exploring Genes of Interest using KBase apps

In order to make RWRtoolkit accessible to users with limited bioinformatic experience, we developed two RWRtoolkit applications within KBase [15] to allow users to input Arabidopsis gene sets. Using the code base of *RWR_make_multiplex*, we assembled 9 multiplex network objects (located at https://github.com/dkainer/RWRtoolkit-data), which were imported into KBase. We first built a KBase application to explore using the *RWR_CV* function of RWRtoolkit (*Find Gene Set Interconnectivity using Cross Validation with RWRtools CV*). We then developed the next application using the *RWR_LOE* function of RWRtoolkit (*Find Functional Context using Lines of Evidence with RWRtools LOE*) to allow users to explore functional Arabidopsis gene-gene linkages from a user’s gene set based on multiple lines of evidence from a random walk exploration of genes within a multiplex network starting from this gene set. Finally, we used *RWR_LOE* embedded within the app to provide a rank-ordered list of genes explored from the user’s gene set.

### 5.5 Exploring Gene Edits with *RWR_LOE*

#### 5.5.1 : Differential Ranking Between Two Gene Sets

We ran *RWR_LOE* two separate times: first with a seed gene set containing *FAE1* (AT4G34520) and *ROD1* (AT3G15820), then with a seed gene set containing *FAE1* and *FAD2* (AT3G12120). Each individual run produced rankings for all 26,605 genes in the multiplex network. We extracted the top 200 ranked genes from running RWR_ LOE on the *FAE1/ROD1* gene pair, and contrasted those to the ranks obtained for those 200 genes when running *RWR_LOE* on the *FAE1/FAD2* gene pair.

#### 5.5.2 : Cytoscape Set Differential

As in the differential ranking analysis, *RWR_LOE* was run using seeds *FAE1* and *FAD2*, and then using *FAE1* and *ROD1* as seeds for a separate ranking analysis. Here, the subnetworks containing the seeds and the top 200 ranked genes were extracted by using the cyto parameter (--cyto 200), generating two separate networks in Cytoscape, named *FAE1.FAD2*, and *FAE1.ROD1*. By subtracting the *FAE1.ROD1* network from the FAE1.FAD2 network using the difference method in Cytoscape, we obtained edges unique only to the *FAE1/FAD2 RWR_LOE* rankings, resulting in the *FAD2* specific network. Conversely, by subtracting the *FAE1.FAD2* network from the *FAE1.ROD1* network, we obtained edges unique only to *FAE1/ROD1 RWR_LOE* rankings, resulting in a *ROD1* specific network. Nodes shared by both networks were obtained via the intersect method in Cytoscape, creating the intersection subnetwork.

#### 5.5.3 : GO Enrichments

Gene set enrichment was run to assess biological functionality for all three distinct subnetworks using ClueGO, obtained from the Cytoscape App Store [50,51]. For each individual network (*FAD2* and *ROD1* specific networks and the intersection network), all gene nodes within each network were loaded into the Load Marker List(s) section and enriched using ClueGO in Functional Analysis mode within the Load Marker List section. The GO Biological Process and KEGG Ongologies/Pathways were selected within the ClueGO settings.

### 5.6 : RWR Shortest Paths

To extract shortest paths between the source genes (*FAE1*, *FAD2*) and the target gene (*EMB3113*), we ran *RWR_ShortestPaths* supplying the source gene set and the target gene set with the cyto parameter as true (--cyto TRUE). The output file contains an edge list with additional metadata for each edge, comprising the shortest paths from all nodes in the source gene set to all nodes in the target gene set.

## Availability of source code and requirements

- RWRtoolkit Source Code available at: https://github.com/dkainer/RWRtoolkit
- *RWR_LOE* and *RWR_CV* are available as web applications (Find Functional Context using Lines of Evidence with RWRtools LOE and Find Gene Set Interconnectivity using Cross Validation with RWRtools CV) through KBase, found at https://www.kbase.us/

## Data Availability

- Pre-Assembled Arabidopsis Networks are available at: https://github.com/dkainer/RWRtoolkit-data.
- Well-Watered Shoot Biomass GWAS Results can be found in Table 3.
- KBase Narrative is publicly available at: https://narrative.kbase.us/narrative/165213.

## Authors’ contributions

David Kainer: Conceptualization, analysis, code, manuscript writing and editing

Matthew Lane, and Kyle A. Sullivan wrote software, analyzed outputs, wrote and edited the manuscript.

J. Izaak Miller and Mikaela Cashman wrote and tested software.

D. Dakota Blair and AJ Ireland created software for the interactive KBase applications.

Mallory Morgan provided manuscript writing and primary analysis.

Ashley Cliff, Jonathon Romero, Angelica Walker, Anna Furches, and Jaclyn Noshay curated data and built networks for multiplex construction.

Hari Chhetri contributed to data collection and analysis by running GWAS on Switchgrass well-watered shoot biomass data.

Yongqin Wang collected phenotypic and genotypic data for Switchgrass well-watered shoot biomass.

Mirko Pavicic contributed to primary manuscript writing and analysis.

Meghan Drake provided project management and manuscript editing.

Natalie Landry acted as project manager for the KBase applications.

Ali Missaoui and Yun Kang supervised collection of phenotypic and genotypic data and provided funding.

Paramvir Dehal and Shane Canon supervised KBase application development.

Daniel Jacobson Conceptualization, funding, supervision, and manuscript editing.

## Acknowledgements

This manuscript has been authored by UT-Battelle, LLC under Contract No. DE-AC05-00OR22725 with the U.S. Department of Energy. The United States Government retains and the publisher, by accepting the article for publication, acknowledges that the United States Government retains a non-exclusive, paid-up, irrevocable, world-wide license to publish or reproduce the published form of this manuscript, or allow others to do so, for United States Government purposes. The Department of Energy will provide public access to these results of federally sponsored research in accordance with the DOE Public Access Plan (http://energy.gov/downloads/doe-public-access-plan). This research used resources of the Oak Ridge Leadership Computing Facility at the Oak Ridge National Laboratory, which is supported by the Office of Science of the U.S. Department of Energy under Contract No. DE-AC05-00OR22725. The switchgrass resequencing data were produced by the US Department of Energy Joint Genome Institute (https://ror.org/04xm1d337; operated under Contract No. DE-AC02-05CH11231) in collaboration with the user community. Sujan Mamidi performed sequence alignments and called the SNP variants, and Jeremy Schmutz leads the HudsonAlpha sequencing efforts.

