## Supplemental Information for "RWRtoolkit: multi-omic network analysis using random walks on multiplex networks in any species"

### SUPPLEMENTAL MATERIAL

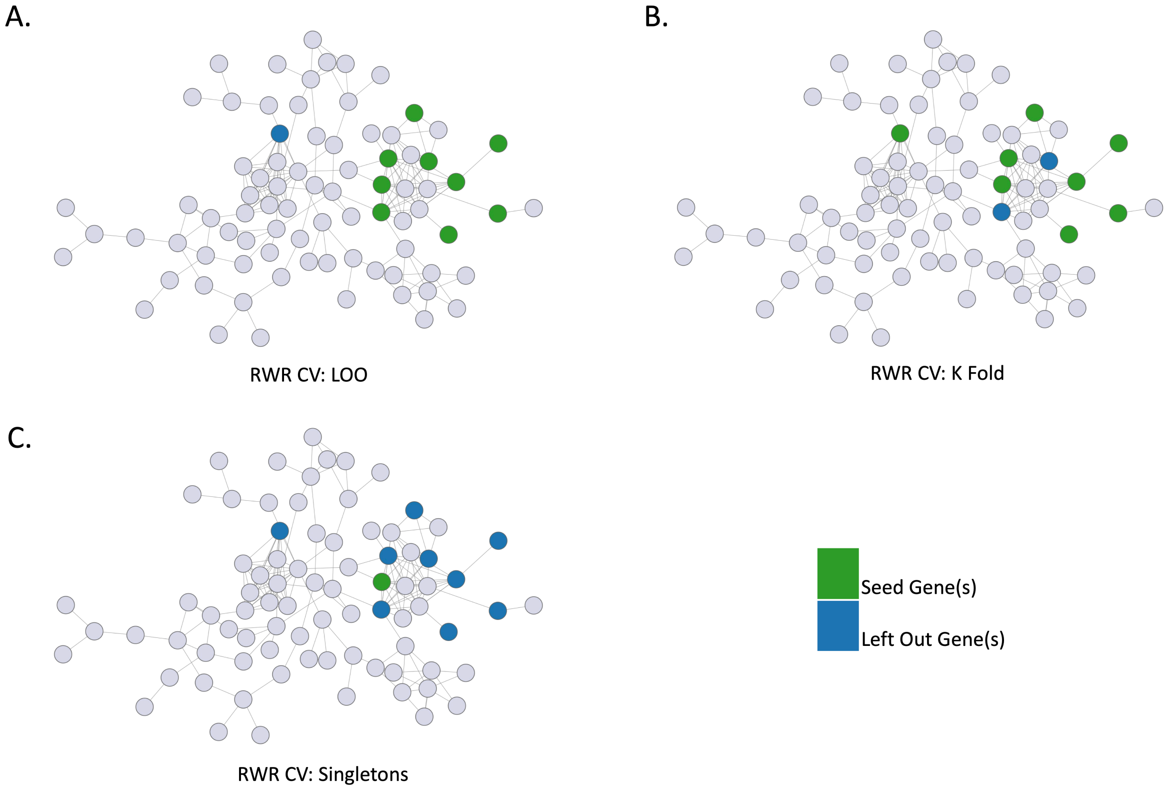


**Supplementary Figure 1. Differences among the methods of *RWR_CV*.** A. Leave One Out (LOO): For each gene within the gene set, LOO uses that gene as a target with the remaining genes in the set used as seeds. B: K Fold: For the total number of folds within the set, N genes are divided by K folds, and randomly chunked without replacement into target sets of size K / N. C. Singletons: For each gene within the gene set, Singletons uses that gene as a seed gene with the remaining genes in the set used as targets.

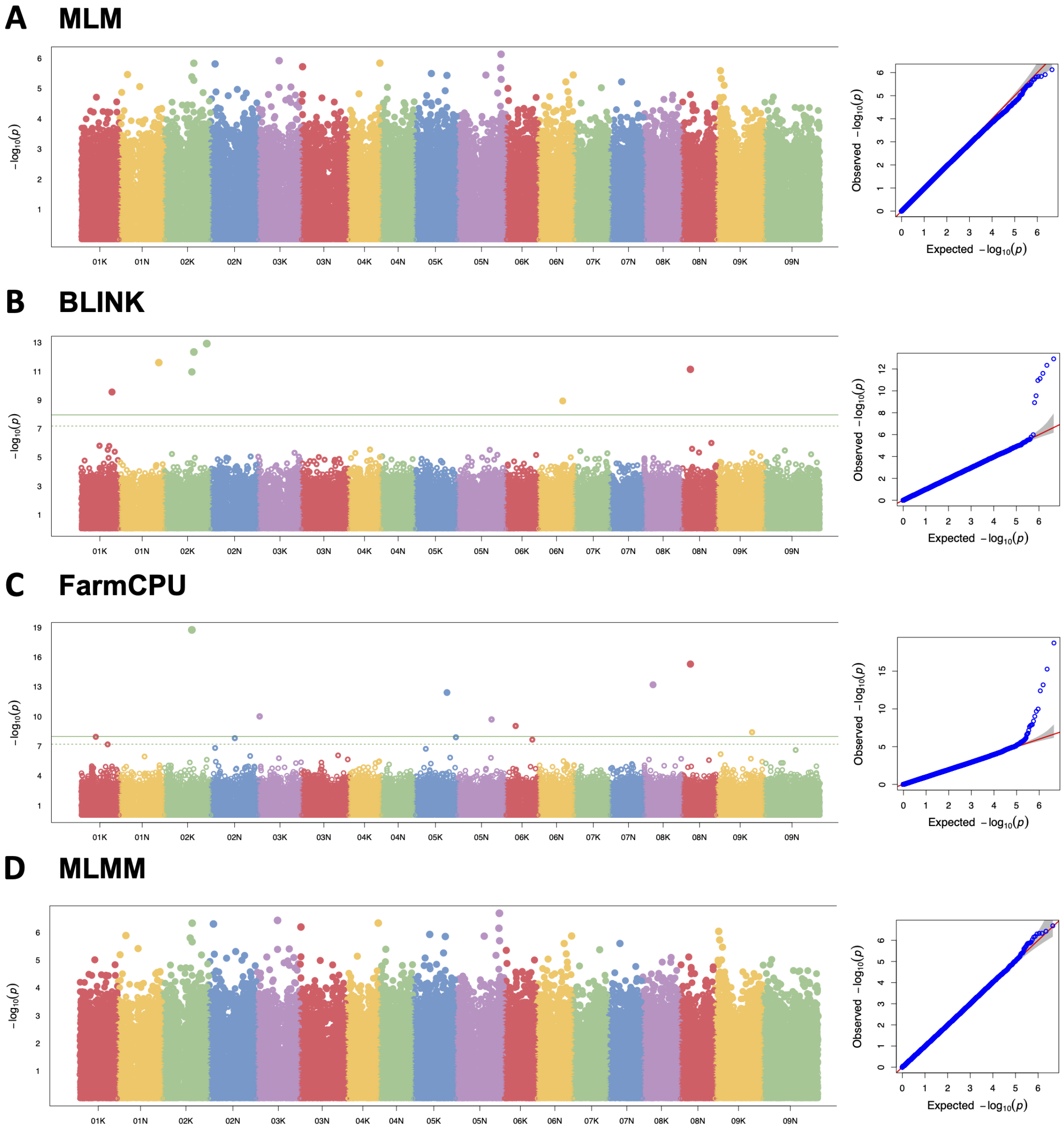


**Supplementary Figure 2.** Switchgrass GWAS Manhattan plots from one single locus (**A**; mixed linear model; MLM) and three multi-locus (BLINK, **B**; FarmCPU, **C**; and MLMM, **D**) using GAPIT. Solid line indicates significance at FDR < 0.2, dotted line indicates suggestive significance. Insets: Q-Q plots for each method.

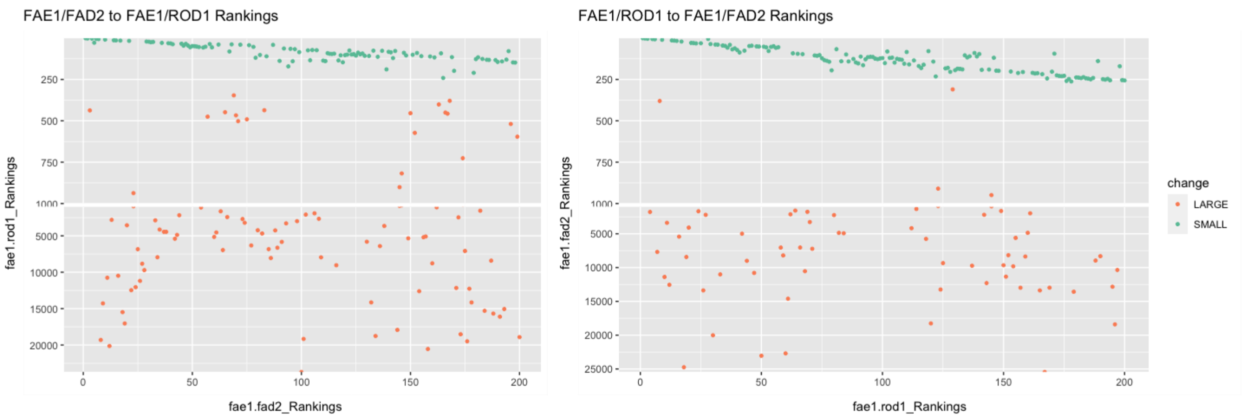


**Supplementary Figure 3. Illustrating *RWR_LOE* Rank Differential between pennycress dual knockout experiments FAE1/FAD2 and FAE1/ROD1.**

The top 200 ranked genes from *RWR_LOE* of FAE1/FAD2 (x-axis, left) and FAE1/ROD1 (x-axis, right) are shown plotted against their corresponding ranks (y-axis) from *RWR_LOE* of FAE1/ROD1 and FAE1/FAD2, respectively. Lower numbers mean better ranking. The top 200 genes for one knockout pair generally achieve a similar ranking in the other knockout pair (blue dots). However, within each experiment, there exist genes that rank far worse when compared to their corresponding rank in the other experiment (orange dots).

| | |
|---|---|
| Gene A | Gene C |
| Gene A | Gene E |
| Gene B | Gene C |
| ... | ... |
| Gene E | Gene D |

**Supplementary Table 1. Example Edge List**

Example network layer edge list with no edge weights to be used as input.

| | | |
|---|---|---|
| Gene A | Gene C | 0.8 |
| Gene A | Gene E | 0.2 |
| Gene B | Gene C | 0.15 |
| ... | ... | ... |
| Gene E | Gene D | 0.99 |

**Supplementary Table 2. Example Weighted Edge List**

Example network layer edge list with edge weights to be used as input.

| | |
|---|---|
| /path/to/layer1 | layer1_name |
| /path/to/layer2 | layer2_name |

| | |
|---|---|
| /path/to/layer3 | layer3_name |
| … | … |
| /path/to/layerN | layerN_name |

**Supplementary Table 3**. **Example Flist**
Example flist in which each row has a full path to an edge list and an associated layer name. While relative paths within the flist can work with respect to the directory from which the program is run, it is highly recommended that users add full paths within their flists.

| network_name | number_of_nodes | number_of_edges | diameter |
|---|---|---|---|
| network_1 | 31 | 98 | 4.863E+00 |
| network_2 | 31 | 98 | 3.710E+00 |
| automated_textmining | 31 | 160 | 1.792E+00 |
| coexpression | 31 | 154 | 5.813E-01 |
| combined_score | 31 | 162 | 2.899E+00 |
| database_annotated | 31 | 136 | 2.889E+00 |
| experimentally_determined_interaction | 31 | 71 | 1.499E+00 |
| gene_fusion | 31 | 5 | 1.809E+00 |
| homology | 31 | 13 | 1.945E+00 |
| neighborhood_on_chromosome | 31 | 87 | 1.848E+00 |

| | | | |
|---|---|---|---|
| phylogenetic_cooccurrence | 31 | 20 | 1.880E+00 |

**Supplementary Table 4. *RWR_netstats* Basic Statistics Example Output**
In the above table, basic statistics are given for all layers of the supplied multiplex as well as the two additional networks supplied to the function.

| | automated textmining | coexpression | combined score | database annotated | experimentally determined interaction | gene fusion | homology | neighborhood on chromosome | phylogenetic cooccurrence |
|---|---|---|---|---|---|---|---|---|---|
| automated textmining | 1.000E+00 | 9.383E-01 | 9.877E-01 | 8.385E-01 | 4.348E-01 | 3.125E-02 | 8.125E-02 | 5.438E-01 | 1.250E-01 |
| coexpression | 9.383E-01 | 1.000E+00 | 9.506E-01 | 7.901E-01 | 3.975E-01 | 2.581E-02 | 3.727E-02 | 5.548E-01 | 8.075E-02 |
| combined score | 9.877E-01 | 9.506E-01 | 1.000E+00 | 8.395E-01 | 4.383E-01 | 3.086E-02 | 8.025E-02 | 5.370E-01 | 1.235E-01 |
| database annotated | 8.385E-01 | 7.901E-01 | 8.395E-01 | 1.000E+00 | 3.019E-01 | 3.676E-02 | 9.559E-02 | 4.204E-01 | 1.223E-01 |
| experimentally determined interaction | 4.348E-01 | 3.975E-01 | 4.383E-01 | 3.019E-01 | 1.000E+00 | 4.110E-02 | 1.507E-01 | 4.107E-01 | 1.974E-01 |
| gene fusion | 3.125E-02 | 2.581E-02 | 3.086E-02 | 3.676E-02 | 4.110E-02 | 1.000E+00 | 0.000E+00 | 5.747E-02 | 8.696E-02 |
| homology | 8.125E-02 | 3.727E-02 | 8.025E-02 | 9.559E-02 | 1.507E-01 | 0.000E+00 | 1.000E+00 | 2.041E-02 | 6.500E-01 |
| neighborhood on chromosome | 5.438E-01 | 5.548E-01 | 5.370E-01 | 4.204E-01 | 4.107E-01 | 5.747E-02 | 2.041E-02 | 1.000E+00 | 9.184E-02 |
| phylogenetic cooccurrence | 1.250E-01 | 8.075E-02 | 1.235E-01 | 1.223E-01 | 1.974E-01 | 8.696E-02 | 6.500E-01 | 9.184E-02 | 1.000E+00 |

**Supplementary Table 5. *RWR_netstats* Pairwise Comparison Between Multiplex Object Layers using Jaccard Similarity Score Example Output**

The above table illustrates Jaccard similarity between each layer in the multiplex.

| | automated textmining | coexpression | combined score | database annotated | experimentally determined interaction | gene fusion | homology | neighborhood on chromosome | phylogenetic cooccurrence |
|---|---|---|---|---|---|---|---|---|---|
| automated textmining | 6.474E-01 | 6.367E-01 | 6.395E-01 | 6.183E-01 | 7.178E-01 | 8.885E-01 | 7.788E-01 | 7.402E-01 | 8.200E-01 |
| coexpression | 2.407E-01 | 2.510E-01 | 2.386E-01 | 1.818E-01 | 3.292E-01 | 4.024E-01 | 4.705E-02 | 3.571E-01 | 2.417E-01 |
| combined score | 9.549E-01 | 9.542E-01 | 9.548E-01 | 9.549E-01 | 9.628E-01 | 9.976E-01 | 9.671E-01 | 9.643E-01 | 9.758E-01 |
| database annotated | 7.969E-01 | 7.908E-01 | 7.933E-01 | 9.449E-01 | 6.160E-01 | 1.000E+00 | 8.855E-01 | 7.216E-01 | 7.756E-01 |
| experimentally determined interaction | 1.685E-01 | 1.433E-01 | 1.726E-01 | 1.341E-01 | 3.938E-01 | 2.822E-01 | 7.429E-01 | 1.582E-01 | 5.923E-01 |
| gene fusion | 1.269E-02 | 6.692E-03 | 1.253E-02 | 1.493E-02 | 3.123E-03 | 4.061E-01 | 0.000E+00 | 2.334E-02 | 4.966E-02 |
| homology | 7.738E-02 | 3.564E-02 | 7.643E-02 | 9.104E-02 | 1.530E-01 | 0.000E+00 | 9.524E-01 | 1.743E-02 | 6.190E-01 |
| neighborhood on chromosome | 2.413E-01 | 2.454E-01 | 2.383E-01 | 2.025E-01 | 2.948E-01 | 5.791E-01 | 4.704E-02 | 4.437E-01 | 2.205E-01 |
| phylogenetic cooccurrence | 1.085E-01 | 6.890E-02 | 1.071E-01 | 1.120E-01 | 1.882E-01 | 2.053E-01 | 9.573E-01 | 7.618E-02 | 8.677E-01 |

**Supplementary Table 6. *RWR_netstats* Pairwise Comparison Between Multiplex Object Layers using Overlap Score Example Output**

The above table illustrates the overlap score between each layer in the multiplex.

| | jaccard |
|---|---|
| automated_textmining | 5.782E-01 |
| coexpression | 5.948E-01 |
| combined_score | 5.771E-01 |
| database_annotated | 5.826E-01 |
| experimentally_determined_interaction | 5.128E-01 |
| gene_fusion | 3.992E-01 |
| homology | 3.701E-01 |
| neighborhood_on_chromosome | 5.866E-01 |
| phylogenetic_cooccurrence | 3.884E-01 |

**Supplementary Table 7. RWR Netstats Multiplex Layers to Ref Net Jaccard example file output.** This file contains statistics for the jaccard score between the total number of edges within the intersection divided by the total number of edges within the union of both networks.

| | overlap |
|---|---|
| automated_textmining | 5.782E-01 |
| coexpression | 5.948E-01 |
| combined_score | 5.771E-01 |
| database_annotated | 5.826E-01 |
| experimentally_determined_interaction | 5.128E-01 |
| gene_fusion | 3.992E-01 |
| homology | 3.701E-01 |

| | |
|---|---|
| neighborhood_on_chromosome | 5.866E-01 |
| phylogenetic_cooccurrence | 3.884E-01 |

**Supplementary Table 8. RWR Netstats Multiplex Layers to Ref Net overlap example file output.** This file sums the edge weights of all intersecting values and then divides those summed edge weights by the total number of edges within the network layer (note: this will be the same as the Jaccard method if all edges are weighted as 1 and the network layer contains all of the edges found within the reference network).

| jaccard | overlap |
|---|---|
| 4.307E-01 | 5.760E-01 |

**Supplementary Table 9. RWR Netstats net to net similarity example file output.** This function compares and calculates metrics for two provided network layers.

| | calculated_tau |
|---|---|
| automated_textmining | 1.134E+00 |
| coexpression | 1.166E+00 |
| combined_score | 1.132E+00 |
| database_annotated | 1.142E+00 |
| experimentally_determined_interaction | 1.006E+00 |
| gene_fusion | 7.828E-01 |
| homology | 7.257E-01 |
| neighborhood_on_chromosome | 1.150E+00 |
| phylogenetic_cooccurrence | 7.615E-01 |

**Supplementary Table 10. RWR Netstats calculated tau example file output.**
This function offers a more advanced method for users to calculate a tau parameter where the parameter is calculated with respect to "overlap" score with respect to a reference network for each layer.

| n_layers | pct_found |
|---|---|
| 1 | 1.85% |
| 2 | 90.12% |
| 3 | 8.02% |
| 4 | 0.00% |
| 5 | 0.00% |
| 6 | 0.00% |
| 7 | 0.00% |
| 8 | 0.00% |
| 9 | 0.00% |

**Supplementary Table 11. RWR Netstats calculate exclusivity file output.**
The table illustrates the overall percentage of edges in the multiplex that exist in N layers. In the above output example, ~2% of edges exist in only 1 layer, ~90% of edges in the multiplex exist in 2 layers, and ~8% of edges exist in 3 layers, with no other edges existing in more than 3 layers.

| NodeNames | Score | rank | num_in_network | num_seeds | networks | modname | seed_geneset |
|---|---|---|---|---|---|---|---|
| H6PD | 7.084E-04 | 1 | 12 | 12 | automated_t...nce | default | setA1 |

| | | | | | | | |
|---|---|---|---|---|---|---|---|
| GPI | 7.036E-04 | 2 | 12 | 12 | automated_t...nce | default | setA1 |
| PFKP | 5.773E-04 | 3 | 12 | 12 | automated_t...nce | default | setA1 |
| HK1 | 5.156E-04 | 4 | 12 | 12 | automated_t...nce | default | setA1 |
| GFPT2 | 4.941E-04 | 5 | 12 | 12 | automated_t...nce | default | setA1 |
| ... | ... | ... | ... | ... | ... | ... | ... |
| GNPNAT1 | 1.826E-04 | 16 | 12 | 12 | automated_t...nce | default | setA1 |
| PGLS | 1.515E-04 | 17 | 12 | 12 | automated_t...nce | default | setA1 |
| KHK | 3.312E-05 | 18 | 12 | 12 | automated_t...nce | default | setA1 |
| IDNK | 1.559E-05 | 19 | 12 | 12 | automated_t...nce | default | setA1 |

**Supplementary Table 12.** ***RWR_LOE*** **output example file output.**
This file denotes node name, associated score from the random walker, rank based off of the associated score, number of seeds in the supplied gene set within the network, the total number of seeds supplied, the multiplex network name (i.e., the combined names of all layers within the network), the modname (alias for experiment name output), and the name of the seed gene set.

| NodeNames | Score | rank | InValset | num_in_network | num_seeds | num_leftout | networks | fold | modname | geneset | seed | leftout | method |
|---|---|---|---|---|---|---|---|---|---|---|---|---|---|
| ENO1 | 1.162E-03 | 1 | 1 | 12 | 8 | 4 | aut...nce | 1 | default | setA1 | many | many | kfold |
| PFKP | 8.004E-04 | 2 | 0 | 12 | 8 | 4 | aut...nce | 1 | default | setA1 | many | many | kfold |

| | | | | | | | | | | | | | |
|---|---|---|---|---|---|---|---|---|---|---|---|---|---|
| GPI | 7.388E-04 | 3 | 0 | 12 | 8 | 4 | aut...nce | 1 | default | setA1 | many | many | kfold |
| ... | ... | ... | ... | ... | ... | ... | ... | ... | ... | ... | ... | ... | ... |
| GFPT1 | 6.626E-05 | 20 | 1 | 12 | 8 | 4 | aut...nce | 3 | default | setA1 | many | many | kfold |
| GFPT2 | 6.518E-05 | 21 | 0 | 12 | 8 | 4 | aut...nce | 3 | default | setA1 | many | many | kfold |
| KHK | 4.215E-05 | 22 | 0 | 12 | 8 | 4 | aut...nce | 3 | default | setA1 | many | many | kfold |
| IDNK | 1.977E-05 | 23 | 0 | 12 | 8 | 4 | aut...nce | 3 | default | setA1 | many | many | kfold |

**Supplementary Table 13.** ***RWR_CV*** **Full Ranks example file output.**
This file contains the output ranking for each *RWR_CV* fold, including the base statistics from *RWR_LOE* seen previously, but also contains InValset, denoting whether the ranked gene was in the target set, fold, number of seed genes, number left out genes, and the method for cross validation.

| NodeNames | meanrank | rerank | InValset | geneset | num_in_network |
|---|---|---|---|---|---|
| ENO1 | 1 | 1 | 1 | setA1 | 12 |
| TPI1 | 1 | 1 | 1 | setA1 | 12 |
| ENO3 | 3 | 3 | 1 | setA1 | 12 |
| GPI | 3.67 | 4 | 0 | setA1 | 12 |
| ... | ... | ... | ... | ... | ... |
| HK3 | 10.67 | 16 | 0 | setA1 | 12 |
| G6PD | 11 | 17 | 1 | setA1 | 12 |

| | | | | | |
|---|---|---|---|---|---|
| PGM1 | 11.67 | 18 | 0 | setA1 | 12 |
| HK2 | 14.67 | 19 | 0 | setA1 | 12 |
| ... | ... | ... | ... | ... | ... |

**Supplementary Table 14.** ***RWR_CV*** **Mean Ranks example file output.**
This file contains the mean ranks for each gene across all folds of *RWR_CV* and reranks each gene with respect to those floating point mean ranks.

| NodeNames | meanrank | rerank | InValset | geneset | num_in_network | TP | FP | cum_TP | cum_FP | FPR | PREC | REC |
|---|---|---|---|---|---|---|---|---|---|---|---|---|
| ENO1 | 1 | 1 | 1 | setA1 | 12 | 1 | 0 | 1 | 0 | 0 | 1 | 0.083 |
| TPI1 | 1 | 1 | 1 | setA1 | 12 | 1 | 0 | 2 | 0 | 0 | 1 | 0.167 |
| ENO3 | 3 | 3 | 1 | setA1 | 12 | 1 | 0 | 3 | 0 | 0 | 1 | 0.25 |
| GPI | 3.67 | 4 | 0 | setA1 | 12 | 0 | 1 | 3 | 1 | 0.053 | 0.75 | 0.25 |
| ... | ... | ... | ... | ... | ... | ... | ... | ... | ... | ... | ... | ... |
| PMM1 | 20 | 26 | 1 | setA1 | 12 | 1 | 0 | 11 | 17 | 0.895 | 0.393 | 0.917 |
| GCK | 21 | 29 | 1 | setA1 | 12 | 1 | 0 | 12 | 17 | 0.895 | 0.414 | 1 |
| KHK | 22 | 30 | 0 | setA1 | 12 | 0 | 1 | 12 | 18 | 0.947 | 0.4 | 1 |
| IDNK | 23 | 31 | 0 | setA1 | 12 | 0 | 1 | 12 | 19 | 1 | 0.387 | 1 |

**Supplementary Table 15.** ***RWR_CV*** **Metrics example file output.**

This file contains the mean ranks denoted above, but additionally does contain metrics such as True Positive (TP), which recalls a gene within the target gene set, False Positive (FP), which recalls a gene not within the target gene set, Cumulative True Positive (cum_TP), which sums all TP as ranks increase, Cumulative False Positive (cum_FP), which sums all FP as ranks increase, False Positive Rate (FPR) which divides cum_FP by the sum of cum_FP and the total number of genes in the network not in the seed set, Precision (PREC) which is cum_TP divided by the sum of cum_TP and cum_FP, and Recall (REC), which is the cum_TP divided by the sum of cum_TP and the difference of cum_TP and num_in_network (False Negative).

| fold | value | measure | geneset |
|---|---|---|---|
| 1 | 0.25 | P@NumLeftOut | setA1 |
| 2 | 0 | P@NumLeftOut | setA1 |
| 3 | 0.5 | P@NumLeftOut | setA1 |
| 1 | 0.398 | AvgPrec | setA1 |
| 2 | 0.162 | AvgPrec | setA1 |
| 3 | 0.514 | AvgPrec | setA1 |
| 1 | 0.249 | AUPRC | setA1 |
| 2 | 0.137 | AUPRC | setA1 |
| 3 | 0.364 | AUPRC | setA1 |
| 1 | 0.174 | ExpectedAUPRC | setA1 |
| 2 | 0.174 | ExpectedAUPRC | setA1 |
| 3 | 0.174 | ExpectedAUPRC | setA1 |
| 1 | 0.500 | AUROC | setA1 |

| | | | |
|---|---|---|---|
| 2 | 0.402 | AUROC | setA1 |
| 3 | 0.609 | AUROC | setA1 |
| meanrank | 0.534 | AvgPrec | setA1 |
| meanrank | 0.475 | AUPRC | setA1 |
| meanrank | 0.486 | AUROC | setA1 |

**Supplementary Table 16.** ***RWR_CV*** **Summary example file output.**
This file contains overall summary statistics for each individual fold as well as mean rank including Average Precision, Area Under the Precision Recall Curve (AUPRC), Expected AUPRC, Area under the Receiver Operating Curve (AUROC).

| from | to | weight | type | weightnorm | pathname | pathlength | pathelements |
|---|---|---|---|---|---|---|---|
| G6PD | HK2 | 8.889E-01 | database_annotated | 6.917E-03 | G6PD_HK2 | 2 | G6PD->HK2 |
| G6PD | HK2 | 9.560E-01 | combined_score | 6.181E-03 | G6PD_HK2 | 2 | G6PD->HK2 |
| ... | ... | ... | ... | ... | ... | ... | ... |
| GFPT1 | HK3 | 1.000E+00 | database_annotated | 7.781E-03 | GFPT1_HK3 | 2 | GFPT1->HK3 |
| GFPT1 | HK3 | 9.259E-01 | combined_score | 5.986E-03 | GFPT1_HK3 | 2 | GFPT1->HK3 |
| ... | ... | ... | ... | ... | ... | ... | ... |

| | | | | | | | |
|---|---|---|---|---|---|---|---|
| MPI | PM<br>M2 | 1.182E-<br>01 | coexpression | 3.058E-<br>03 | PKLR_PMM2 | 4 | PKLR->GPI-<br>>MPI->PMM2 |
| MPI | PM<br>M2 | 8.968E-<br>01 | automated_textmining | 8.657E-<br>03 | PKLR_PMM2 | 4 | PKLR->GPI-<br>>MPI->PMM2 |

**Supplementary Table 17: *RWR_ShortestPaths* example file output.**
This file extracts all connections within a path between nodes in the source gene set and target gene set. Edges for each path are saved with the layers in which those edges exist as well as each edge's original edge weight, normalized edge weight, a unique path name id, total path length, and a list of all elements within the path.

| | |
|---|---|
| setA | gene_A |
| setA | gene_B |
| setA | gene_C |

**Supplementary Table 18: RWR Gene Set Example Format**
Most RWRtoolkit commands require the user to input a set of genes of interest, known as a gene set. A gene set is a tab-delimited file where the first column is the name of the gene set itself, the second column is the unique gene IDs, and the third (optional) column is a weight per gene which can be used to influence the random walk if desired. The genes in the gene set can be used as seeds for random walks, target genes in *RWR_LOE* and *RWR_ShortestPaths*. An example of a gene list can be seen in Table 4.
